# Mice adapt their learning rate to stochasticity and volatility through a simple heuristic

**DOI:** 10.64898/2026.09.24.754025

**Authors:** Coline Chevallier, Juan Cobos, Emma Debos, Emma Marillat, Julie Perroy, Jérémie Naudé

## Abstract

Learning from outcomes requires weighing each surprise by how much it is likely to mean. Two properties of the environment set that weight in opposite directions: volatility, the rate at which contingencies change, and stochasticity, the noise in outcomes around a stable contingency. Whether animals separate the two is unresolved, in part because varying reward probability in a reversal task makes outcomes noisier and the two options harder to tell apart at once. We trained male and female mice on a probabilistic reversal task for intracranial self-stimulation, scaling reward magnitude with reward probability so that the expected reward of each option and the difference between them stayed constant while outcome variance varied, and crossing this with different reversal rates. Simulations show that adapting the learning rate alone and adapting the decision policy alone are near-equivalent solutions across the environments we tested. Mice adopted the first adjustment: their fitted learning rate rose with volatility and fell with stochasticity, in both directions predicted by reward maximization, while their inverse temperature stayed fixed. This joint dependence was reproduced only by a meta-learning rule that reads stochasticity from the size of prediction errors and volatility from how often recent outcomes favored switching.

## 1 Introduction

Adapting behavior to outcomes is what allows an animal to survive in a changing world [1, 2, 3]. Yet similar surprising outcomes may call for opposite adaptations. Consider an animal that returns to a food source that usually pays off and finds nothing. An isolated failure may reflect nothing more than the probabilistic nature of the source, in which case the animal should ignore it and return. Successive failures may instead signal that the source has stopped paying, in which case it should be abandoned. The same outcome warrants opposite responses, and the animal cannot tell which one applies without an estimate of the kind of uncertainty it faces [4].

Two properties of the environment set whether a surprise is informative. Stochasticity, the noise in outcomes around a stable state, determines how often a good choice goes unrewarded. Volatility, the rate at which that state itself turns over, determines how often contingencies actually change. The two are independent : an environment can be stable but noisy or near-deterministic but changeable. Volatility and stochasticity push the weight a surprise should carry in opposite directions [5, 6]. That weight is the learning rate of a reinforcement learner, an agent that assigns each option a value, chooses on the basis of these values, and revises them in proportion to the departure of the outcome from what was expected [7, 3]. Tuning the learning rate to the joint demands of change and noise is a central problem of learning under uncertainty [8, 9, 10, 11].

Tuning the learning rate to opposite forms of uncertainty is difficult because an animal observes neither quantity directly, yet could arrive at an appropriate learning rate from experience alone. Different classes of model answer this challenge differently. A model with a fixed learning rate answers that an agent behaves differently across environments only because the outcomes differ. Descriptive models, such as recursive logistic regressions, predict the next choice from the recent run of choices and outcomes and recover the graded, history-dependent switching that a fixed rate misses, but only describe the dependence instead of explaining it [12, 13, 14]. Bayesian inference models take the normative problem literally, by considering that agents maintain beliefs over the hidden state of the world, at the cost of an explicit internal model of the task and continuous tracking of latent variables the animal never observes [15, 16, 8]. Such an observer must be handed the state space, the form of the reward distributions or the transition structure. So the question is whether rules of far lower complexity suffice. This is what meta-learning models provide, by letting the learning rate itself become a variable driven up when recent outcomes suggest the world is changing, and down when they suggest it is only noisy, read approximately off the stream of prediction errors [17, 18, 19, 20]. Meta-learning is used here in this sense, an explicit rule that sets a learning parameter from ongoing experience, and not in the sense of a recurrent network trained until a learning algorithm emerges in its activity.

These accounts remain hard to separate on behavior alone. During stable phases they predict very similar choices [21, 22, 23], and when the environment changes they all produce some adaptation. Telling them apart requires a task in which volatility and stochasticity vary independently, and the human studies that crossed the two did not converge. Facing outcomes drawn from a Gaussian whose mean set which option was better and whose standard deviation set outcome noise, participants raised their learning rate with volatility and showed no normative response to noise, and a Bayesian observer analysis indicated that they were adapting to a coarse estimate of noise and attributing it to volatility [24]. A design in which both sources are binary and explicitly described to participants recovers both effects, learning rates rising with volatility and falling with stochasticity [25]. Whether the negative dependence on stochasticity is a robust property of adaptive learning is therefore still open, and it has not been tested in rodents yet.

Testing it in a reversal task first requires resolving a confound. Manipulating noise as the variance of a graded outcome leaves the expected difference between the options untouched. Manipulating it through reward probability, the format available in animal reversal learning, does not: moving both probabilities toward chance makes outcomes noisier and, at the same time, makes the two levers harder to tell apart. We therefore designed a probabilistic reversal learning task [26, 27] in which the two sources of uncertainty are manipulated independently. Volatility was set by the frequency of reversals. Stochasticity was set by moving both reward probabilities toward or away from chance, with reward magnitude scaled so that the expected reward of each option, and the difference between them, stayed constant. Only outcome variance changed, and mice were never instructed about either source. Mice performed the task for intracranial self-stimulation [28, 29, 30, 31], which holds motivation stable across sessions, sets the reward magnitude directly, and yields the thousands of trials required to estimate choice statistics within each environment.

Reward maximization does not by itself dictate which parameter should move. Simulating the task shows that an agent adapting only its learning rate and an agent adapting only its decision policy earn almost the same reward across the whole stochasticity-volatility plane, so the environment requires that one of the two be retuned and leaves open which one [17]. Mice retuned the first. They raised their learning rate with volatility and lowered it with stochasticity, in both directions predicted by the normative analysis, while their inverse temperature stayed fixed across the same environments [32]. We account for this with a meta-learning model in which stochasticity is read from a running average of unsigned prediction errors [8] and volatility from an incompatibility signal that rises whenever repeating a choice stops paying or switching starts paying [33], requiring no inference over hidden states [34]. We have shown in rats that orbitofrontal noradrenaline supports the adjustment of the learning rate to volatility [33]; the incompatibility signal is the computation such a system would have to carry, isolated here in a task that separates volatility from stochasticity. This rule is not the one favored by penalized fit quality, and the discrepancy is itself informative: a descriptive account reaches a lower BIC by absorbing the cross-environment structure into its parameters rather than by computing it. By isolating the dynamic learning rate within a high-throughput behavioral task in mice, this framework brings adaptive learning within reach of its neural dissection.

## 2 Materials and Methods

### 2.1 Experimental model and study design

20 male and 20 female C57BL/6J (Wild-type) from Janvier Labs (France) were used for all the experiment, with 11 animals excluded (final : 15 male mice, 14 female mice). All the mice weighed between 25 and 35 g and were 2 months old upon arrival. The animals were housed in groups of two or three in a room with a health status complying with the European standard established by the Federation of European Laboratory Animal Science Associations, where the temperature was maintained at 20 ° C ± 2 ° C, humidity was controlled, and a 12-hour light/12-hour dark circadian cycle was maintained. All experiments were conducted during the light phase of the cycle. The cages were carefully enriched to accommodate the full range of the mice natural behaviors (hiding, climbing, digging and burrowing, and gnawing). They had continuous access to water and food. The experiments received approval from the ethics and animal experimentation committee of the French Ministry of Higher Education and Innovation (reference number of the project: APAFIS #2209-50949.).

### 2.2 Stereotaxic surgery and implantation

The mice were anesthetized with a mixture of oxygen (1 L/min) and isoflurane (1000 mg/g, Vetflurane - 1–3%), then placed into a stereotaxic frame. We tested the animal’s reflexes to ensure adequate anesthesia, then injected 0.1 mL of lidocaine (Lurocaine^®^) via subcutaneous injection at the skull, followed by 0.1 mL of Buprenorphine (BupreCare ^®^) via intraperitoneal injection. We applied an ophthalmic gel (vitamine A) to prevent eye dryness and use a heating pad to maintain the animal’s normal body temperature. We positioned the animal so that the skull is level, then made a 1.5 cm skin incision. We followed a 5-step antisepsis protocol using povidone-iodine. We then verified that the skull was properly aligned by taking stereotaxic coordinates. Once the zero point has been correctly established at the bregma, we identified the coordinates for electrode implantation. We performed a craniotomy using a drill above the coordinate points (Electrode implantation : ML + or -1.2, AP -1.4, DV -4.8)([35]). We used dual channel stimulation electrodes (MS303/3B/SPC, Protech) twisted stainless steel. The electrode was then implanted unilaterally for inrrcranial self-stimulation (ICSS) in the medial forebrain bundle (MFB) [28, 29, 30, 31], with the implantation side randomly counterbalanced over the experimental group. To fix it to the mouse skull, we used a self-polymerizing adhesive resin cement (Super Bond – SUN MEDICAL). After suturing and recovery of the animal, we carried out regular monitoring over two days by repeatedly assessing the key criteria of the Mouse Grimace Scale (MGS). If necessary, we administered an analgesic (Buprenorphine). A one-week recovery period was allowed between the surgery and the start of the experimental protocol.

### 2.3 Reversal probabilities task

#### 2.3.1 Apparatus

The behavioral task was carried out in an operant behavioral chamber (from MedAssociates equipment) measuring 30 × 24 × 25 cm, made of metal. Two levers spaced 15 cm apart were continuously available. The animals were connected through the stimulation electrode and received rewards in the form of MFB stimulation. The stimulations consisted of square-wave signals calibrated in amplitude and in the number of pulse trains by a pulse generator and an isolated constant-current stimulator. The current amplitude was adjusted individually for each animal during the conditioning phase to ensure a sufficiently motivating reward, stabilizing the number of lever presses between 500 and 1000 per session without inducing compulsive behavior in the animal. After the initial conditioning phase, this value remained fixed for the rest of the experiment. The number of pulse trains, however, was adjusted according to the environment in order to maintain a constant average available reward per choice across all sessions.

The experimental room was maintained at a temperature of 20 ± 2°C, and the light intensity remained constant across sessions and days. Experimental events were controlled by a program using the Med-Soft-IV software, and data recording was also handled by this software.

#### 2.3.2 Task structure

The experiment was conducted in three successive phases, with daily sessions of 30 minutes. Firstly, an initial Conditioning phase in a deterministic environment (7-10 days). During this phase, the animals were trained in an environment which two distinct actions (pressing the left or the right lever) each systematically led to a reward. This initial phase allowed the mice to learn the association between performing an action and receiving a reward. The average reward available for each lever press was 20 ICSS pulses.

The second phase consisted of contingency learning and behavioral flexibility (3 days). The animals were placed in the same deterministic environment, but only one of the two possible actions was associated with a reward. After 100 responses, the reward contingencies were reversed. This alternation was repeated throughout each session to train the animals to adapt their behavior to changes in contingency.

During the third phase (6 days), the animals were placed in uncertain environments whose statistical properties varied across sessions [26, 27, 14].

Two environmental statistics were defined to characterize uncertainty.

Reward stochasticity was defined as the variance around a given mean reward probability associated with each lever. We tested two pairs of reward probabilities for the high-reward lever (and, respectively, the low-reward lever) 80%–20% and 64%–16%. A compensation scheme was implemented under the assumption that the subjective utility of ICSS was linear with the number of stimulation pulses within the individually calibrated range, such that *P U* (*M*) = *P M* (see Fig. 1). This scheme yielded 20 pulses in the 80%–20% condition (M=1) and 25 pulses in the 64%–16% condition (M=1.25). We verified that neither press rates nor inter-press intervals differed systematically between the 80/20 (M=1) and 64/16 (M=1.25) conditions, consistent with matched subjective value. Stimulation waveforms were also verified during behavior using analog recordings together with Med-Soft-IV command logs. Environmental volatility was defined as the number of lever presses over which a lever’s reward distribution remained unchanged (i.e., the block length). We examined three block lengths: 25, 50, and 100 presses. The combination of these parameters yielded six uncertain environments with different levels of stability and stochasticity (Fig. 3). The order of sessions with the different environments was pseudo-randomized following a latin square design over the experimental group. For each animal and each session we recorded, for every lever press, the identity *C_n_* of the chosen lever at trial *n* (Left : *C_n_* ≡ *L*, Right : *C_n_* ≡ *R*), whether it was rewarded (*r*(*n*) = 1) or not (*r*(*n*) = 0), as well as the timestamp of the press.

**Figure 1:**
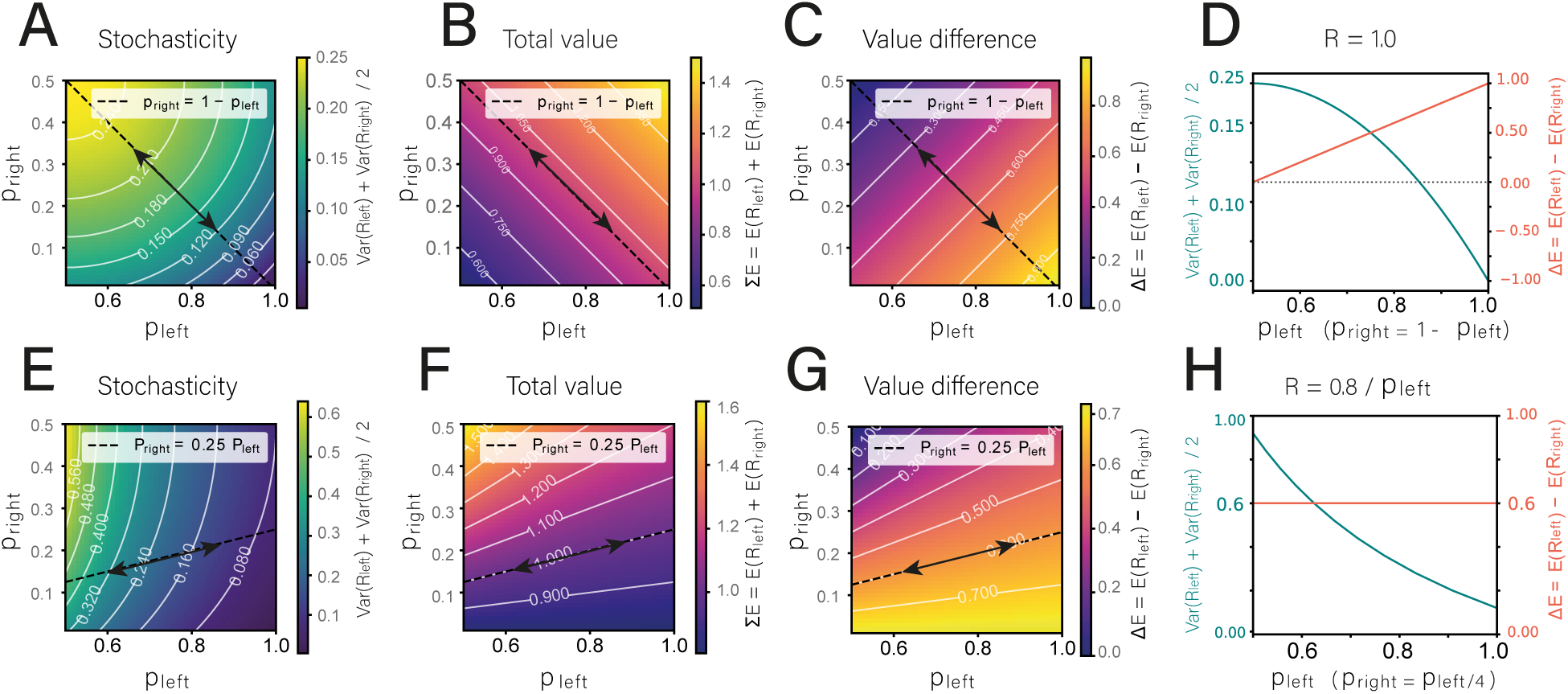
Varying reward probability changes outcome variance and option discriminability at once, unless reward magnitude compensates. **(A–D)** Standard design, in which the two reward probabilities trade off against each other (*p_R_* = 1 − *p_L_*) at a fixed reward magnitude. Maps are drawn over all probability pairs (*p_L_, p_R_*), with contours giving iso-levels of the plotted quantity and the diagonal line marking the design path *p_R_* = 1 − *p_L_*. Moving both probabilities toward chance along that path raises the mean outcome variance of the two options (A), which is the intended manipulation of stochasticity, and leaves the mean available reward unchanged (B), but it also shrinks the difference in expected reward between the options (C), the quantity that sets how discriminable they are. **(D)** The same two quantities plotted against each other along the design path: outcome variance and value difference move together, so a change in behavior across such conditions cannot be assigned to one rather than the other. **(E–H)** Same four quantities under the compensated design, in which reward magnitude is scaled with reward probability so that *P M* is constant for each option; the design path is now *p_R_* = 0.25 *p_L_*. Outcome variance still varies (E), the mean available reward is still fixed (F), and the difference in expected reward is now fixed as well (G), so that outcome variance moves at constant value difference (H). Overlaid points mark the two experimental conditions, 80/20 with *M* = 1 and 64/16 with *M* = 1.25.

### 2.4 Data analysis

#### 2.4.1 Reward statistics of the environments

To characterize how the reward-probability pairs used in the task relate stochasticity to task difficulty, we computed three summary statistics of the reward structure for every pair of reward probabilities (*p_L_, p_R_*), under two designs. In the classical design, each rewarded press delivers a fixed magnitude (*M* = 1) and the two options trade off their probabilities (*p_R_* = 1 − *p_L_*). For a Bernoulli outcome of probability *P* and magnitude *M*, the expected reward of an option is *PM* and its variance is *P* (1 − *P*)*M*^2^. The three statistics, each averaged over the two options, are the mean outcome variance,

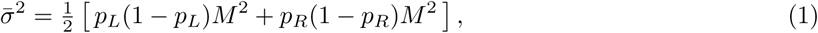

the mean available value 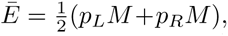, and the value difference between options Δ*E* = | *p_L_M* −*p_R_M* |, which we take as an index of task difficulty (small Δ*E* making the two options hard to tell apart). Varying stochasticity along *p_R_* = 1 − *p_L_* changes the mean variance but also changes Δ*E*, so the two are confounded.

In the magnitude-compensated design, the reward magnitude of each option is set so that its expected value is held constant, *PM* = *c*, i.e. *M* = *c/P*. Each option then delivers the same expected reward regardless of its probability, so *Ē* and Δ*E* no longer vary with the probability pair, while the mean variance 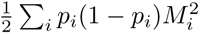 still tracks stochasticity. Plotting these statistics as maps over (*p_L_, p_R_*), with the experimental conditions overlaid, shows that magnitude compensation dissociates outcome variance from value difference, which co-vary in the classical design (Fig. 1).

#### 2.4.2 Inclusion criteria

We included in our analyses all mice that met a predefined set of inclusion criteria, thereby ensuring data quality, reproducibility, and the statistical robustness of the results. Specifically, we retained mice that successfully learned the operant conditioning task, required a stimulation current not exceeding 200 µA, and reached a stable level of responding between 500 and 1,000 lever presses per session by the end of the initial conditioning phase. This led to the exclusion of 6 mice (3 male, 3 female). During phase 3, animals were required to perform between 300 and 1000 lever presses per session. Failure to meet these criteria in a given session resulted in the exclusion of all sessions from the corresponding animal to keep a balanced design. This led to the exclusion of 5 mice (2 male, 3 female).

#### 2.4.3 Analysis

No formal power calculation was performed before the experiment. The design was sized for parameter estimation rather than for a group contrast: each of the 29 mice contributed one session in each of the six environments, with 300 to 1000 lever presses per session, for a total of 144647 scored trials, that is 4988 per animal and 831 per animal and environment on average. What limits the precision of a computational parameter is the number of trials per subject and per condition at least as much as the number of subjects [36], and the recovery analyses of Section 3.4 (Fig. 4E-H) quantify that precision directly for this design rather than by rule of thumb: at these session lengths, *α* and *β* are both recovered together with their dependence on the environment. Sex was not a design factor, and the sex comparisons reported below are exploratory.

*Success rate* is the proportion of lever presses that were rewarded. It was computed for each animal in each of the six experimental environments, defined by the combination of stochasticity level (low stochasticity, LS; high stochasticity, HS) and volatility level (low volatility, LV; medium volatility, MV; high volatility, HV). It measures the reward the animal actually collected. The two measures defined below, perseverative errors and post-reversal adaptation, are instead defined on the identity of the pressed lever, that is on whether the animal chose the lever programmed as better, independently of whether that particular press was rewarded.

*Win–shift and lose–shift* behavior provides a measure of behavioral sensitivity to environmental feedback. For each animal in each of the six environments defined above, we computed the win–shift ratio as the probability of switching to the alternative option following a rewarded trial. Conversely, the lose–shift ratio was computed as the probability of switching following an unrewarded trial. These ratios were calculated independently of whether the previously selected lever was the best one. The standard error of the mean (SEM) was then calculated across animals for each environment.

*Perseverative errors* provide information about the animal’s tendency to perseverate, or conversely, its ability to rapidly detect and adapt to changes in the environment [27, 37]. They were computed as the number of consecutive presses on the worst lever following a contingency reversal, up to the first press on the optimal lever.

*Post-reversal adaptation* provides a complementary measure of the animal’s ability to adapt to contingency reversals. For each contingency reversal, we computed the probability of selecting the best lever for each trial position relative to the reversal. These probabilities were first averaged across reversals for each animal and then averaged across animals. For each environment, we plotted the probability of selecting the best lever as a function of trial position relative to the contingency reversal.

*Probability to shift after a certain pattern of choices* highlights whether specific combinations of three consecutive choices are preferentially expressed in certain environments. We considered all possible combinations of three-choice sequences, including switches (shift, ab), repeated choices (stay, aa), rewarded choices (A, B), and unrewarded choices (a, b). The probability of switching after each triplet was first averaged across sessions for each animal, and then averaged across animals [13].

For all measures, the standard error of the mean (SEM) was calculated across animals for each environment.

### 2.5 Computational modeling

#### 2.5.1 Simple behavioral model

We studied the behavior of mice in a volatile and stochastic environment during a probability reversal learning task. We first used simple behavioral win-stay/lose-shift (WSLS) model which promotes repeating the same choice after a rewarded outcome, while switching to an alternative choice following a non-rewarded outcome.

In our experimental paradigm with a binary choice between two options, the decision rule governs the probability to choose one of the options, hereafter arbitrarily denoted by “L” (for “Left” lever), the other option being denoted by “R” (for “Right” lever).

##### Decision rule

The probability of choosing option L on trial n+1, follow :

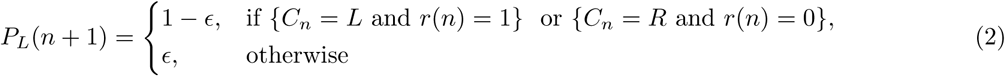

with *ɛ <* 0.5 the single free parameter, i.e. the probability of departing from the win-stay/lose-shift rule. Equivalently, this corresponds to a stay probability after reward *p*_stay_ = 1 − *ɛ* and a shift probability after omission *p*_shift_ = 1 − *ɛ*, so that the model has one degree of freedom.

#### 2.5.2 Reinforcement-learning models

Reinforcement learning models and variants are used as generative models of our mice behavior in the task.

##### Decision rule

This probability on trial n, *P_L_*(*n*), is basically expressed as a function of the difference between values associated with each option at trial n, *V_L_*(*n*) and *V_R_*(*n*). It is formulated as the Softmax rule [3, 21]:

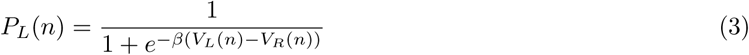

with *β*, the inverse temperature parameter that balances exploration and exploitation. When *β* = 0, *P_L_*(*n*) = 0.5 and the agent makes its choice independently of the options values (pure exploration). As *β* becomes larger and larger, the agent tends to choose the best option with higher and higher probability (favoring exploitation).

As a generalization, an additional choice stickiness parameter *κ* ≥ 0 is introduced to represent the cost due to switching option [38, 39]. It is added to the value associated with the last chosen option. In the case this last option is L, we have:

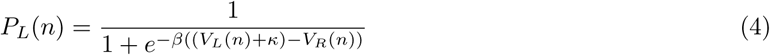

Denoting *C_n_*the option chosen at trial n (L or R), the reward prediction error (RPE) is calculated as:

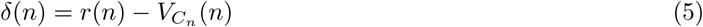

with *r*(*n*) = *M* the reward magnitude if the option was rewarded and *r*(*n*) = 0 otherwise.

Below, a learning rule will update (*V_L_, V_R_*) depending on *δ*(*n*).

##### Classical learning rule

The update of the option values follows the following Rescorla-Wagner rule [7, 3], based on the reward prediction error:

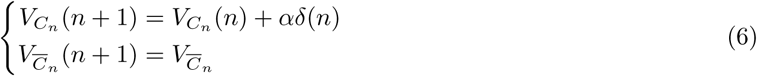

where *α* ∈ [0, 1] is a learning rate parameter and *C⎺_n_* denotes the unchosen option.

Note that first equation in Eq 6 fully reads:

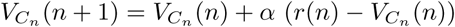

and can be interpreted as a discrete step exponential relaxation towards *r*(*n*).

##### Coupled learning rule

As we considered two-options environment, the values are updated symmetrically [33, 13]. The unchosen lever loses (resp. gains) as much value as the chosen option gains (resp. loses) in the event of a reward (resp. omission).

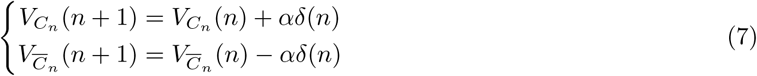

##### Asymmetric learning rule

In this variant [40], the learning rate of the option values depends on whether or not a reward was obtained on trial n.

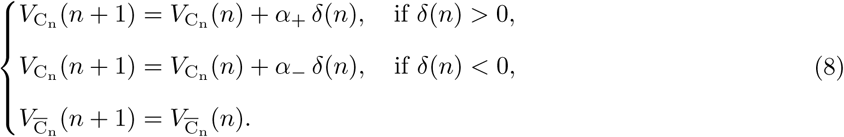

#### 2.5.3 Meta Reinforcement-learning models

##### Meta-RL-surprise

In the meta-RL models [41, 4, 8], the learning rate becomes variable across trials. *α*(*n*) is now a function of estimates of volatility *λ*(*n*) and stochasticity *ɛ*(*n*), following:

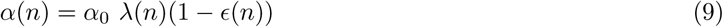

Evolution of *ɛ*(*n*) and *λ*(*n*) are based on reward prediction errors *δ*(*n*) only.

For the stochasticity estimate *ɛ*(*n*), this model considers that animals use the absolute value of the RPE [8, 42, 41], which represents the discrepancy between expectation and values:

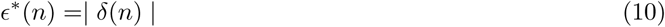

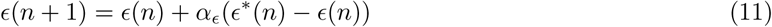

such that if discrepancy occurs often (high stochasticity), *ɛ* increases.

Because *ɛ^∗^*(*n*) =| *δ*(*n*) | is expressed in reward units, the stochasticity estimate *ɛ*, and the term weighting it in the learning rate, carry the scale of the reward magnitude *M*, whereas the learning rate of the fixed-rate models is dimensionless (see Section 3.4). The two magnitudes used here differ by a factor 1.25, so in both meta-RL models part of the difference in learning rate between the two stochasticity conditions is a scale effect on *ɛ* in addition to the change in outcome variance. The two models share this property, and it therefore does not contribute to the difference between them.

For the volatility estimate *λ*(*n*), this model considers that animals monitor the need to adapt the estimation of stochasticity:

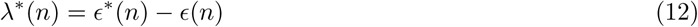

with

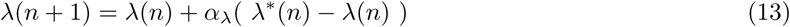

##### Meta-RL-incompatibility

This model is inspired by meta-RL-surprise above, with two modifications.

First, the effective learning rate combines the two uncertainty estimates additively rather than multiplicatively:

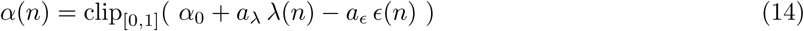

where *a_λ_* and *a_ɛ_* are non-negative gain parameters weighting the volatility and stochasticity estimates, and clip[0,1] bounds the learning rate to [0, 1]. Volatility thus raises the learning rate while stochasticity lowers it, in line with the normative prediction (Fig. 2). This additive form avoids the compensation between *ɛ* and *λ* that arises in the multiplicative coupling, which cancels the volatility signal.

**Figure 2:**
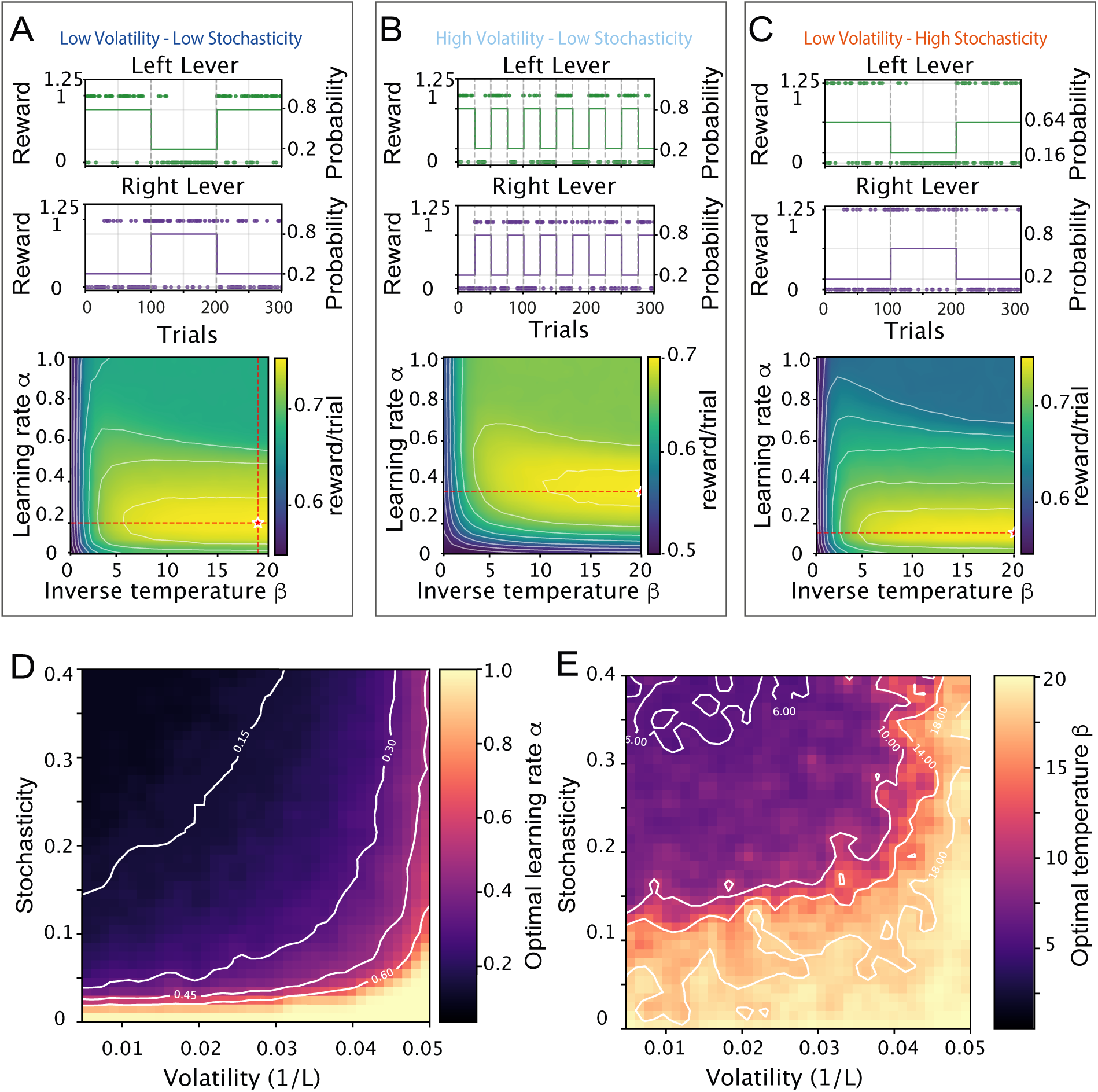
Normative predictions from an optimal reinforcement learning model. **(A–C)** For three example environments, average reward per trial obtained by simulating the RL model with choice stickiness as a function of the learning rate *α* (inverse temperature held fixed) and of the inverse temperature *β* (learning rate held fixed). Examples are given for low stochasticity and low volatility (A); low stochasticity and high volatility (B); and high stochasticity and low volatility (C). The reward landscape is sharply peaked along *α* but shallow along *β*. **(D)** Reward-maximizing learning rate *α*^opt^ across the stochasticity (reward probability) by volatility (1*/L*, block length) plane, with the inverse temperature held fixed (*β* = 2.5). *α*^opt^ increases with volatility and decreases with stochasticity. **(E)** Reward-maximizing inverse temperature *β*^opt^ across the same plane, with the learning rate held fixed (*α* = 0.4). *β*^opt^ increases with volatility and decreases with stochasticity. Averaged over the plane, the regret relative to *R*_max_ = 0.8 reward per trial is 0.094 for the *α*-adapting agent and 0.108 for the *β*-adapting agent.

**Figure 3:**
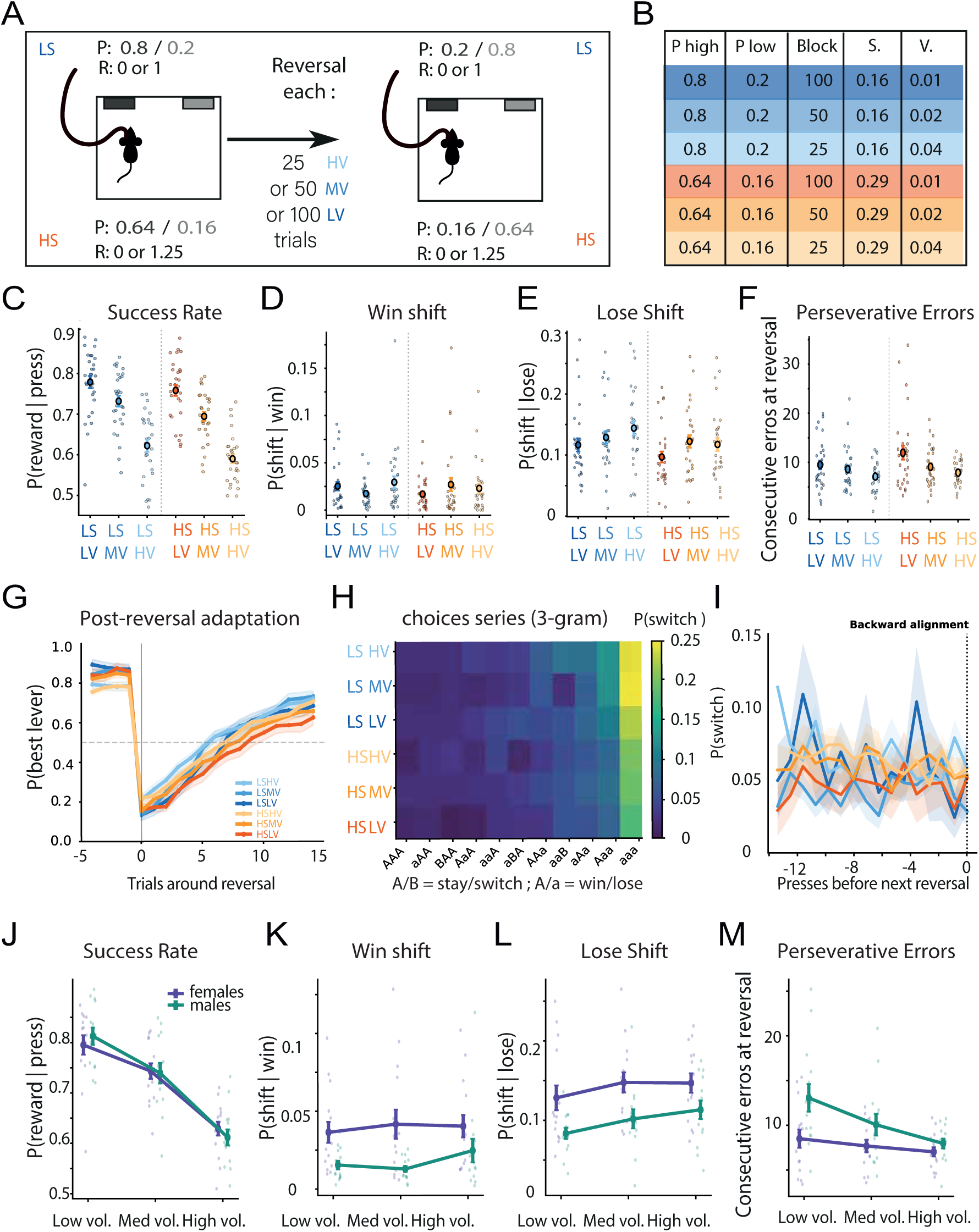
Behavioral signatures of adaptation across the six environments. **(A)** Schematic of the probability reversal task with independent co-variation of volatility and stochasticity. **(B)** The six uncertain environments, defined by the compensated reward-probability pair (stochasticity, S) and the block length between reversals (volatility, V). **(C)** Performance is highest in stable, deterministic environments and declines with both factors (stochasticity *F* (1, 28) = 11.92, *p* = 0.0018; volatility *F* (2, 56) = 78.66, *p <* 10*^−^*^9^; two-way RM ANOVA). **(D)** Win-shift probability is flat across the six environments (stochasticity *F* (1, 28) = 0.21, *p* = 0.65; volatility *F* (2, 56) = 0.83, *p* = 0.44). **(E)** Lose-shift probability is graded by both factors (stochasticity *F* (1, 28) = 7.63, *p* = 0.010; volatility *F* (2, 56) = 4.61, *p* = 0.014; interaction *F* (2, 56) = 1.04, *p* = 0.36). **(F)** Mean number of consecutive perseverative errors following a reversal, depending on volatility (*F* (2, 56) = 7.61, *p* = 0.0012) and, more weakly, on stochasticity (*F* (1, 28) = 4.63, *p* = 0.040; interaction *F* (2, 56) = 1.39, *p* = 0.26). **(G)** Choice of the more rewarded option around a reversal, showing how adaptation accelerates with volatility and decelerates with stochasticity. **(H)** Probability to switch depending on the history of the 3 previous choices (3-gram), with the initial direction in the sequence referred to as A/a and the other direction as B/b, capital letters (A/B) indicating rewarded presses. **(I)** Probability of switching as a function of press position relative to the next reversal. Switching over the last five presses of a block does not differ from the stable plateau (paired *t*(28) = −0.15, *p* = 0.88). **(J)** Success rate did not differ between male and female mice (*F* (1, 27) = 0.007, *p* = 0.9) and displayed a similar dependence on volatility (*F* (2, 54) = 77.8, *p <* 0.001). **(K)** Win-shift was higher in male than female mice (*F* (1, 27) = 9.5, *p* = 0.004) and showed a similar independence from volatility (*F* (2, 54) = 0.8, *p* = 0.44). **(L)** Lose-shift was higher in male than female mice (*F* (1, 27) = 8.4, *p* = 0.004) and displayed a similar dependence on volatility (*F* (2, 54) = 7.9, *p* = 0.009). **(M)** Perseverative errors were more numerous in female than in male mice (*F* (1, 27) = 6.48, *p* = 0.017) and increased with volatility in both sexes (*F* (2, 54) = 8.03, *p <* 0.001).

In the learning equations, the parameter *α* is replaced by a variable, denoted by *α*(*n*), as defined above. Function *ɛ*(*n*) is left unchanged:

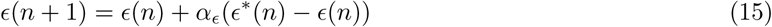

with *ɛ^∗^*(*n*) given by Eq 10.

Second, for the volatility estimate *λ*(*n*), animals are assumed to track an “incompatibility” signal that reads the sign of the choice-conditioned outcome. Writing the signed outcome *u*(*n*) = 2 *r*(*n*) − 1 ∈ {−1, +1} and *s*(*n*) = 1(*C*(*n*) ≠ *C*(*n* − 1)) for a repeated choice, the incompatibility target is:

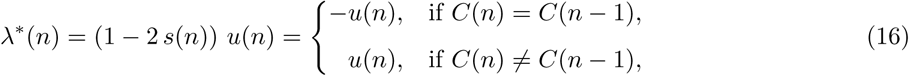

Thus, repeating a choice and receiving no reward, or switching choices and receiving a reward, signals an incompatibility between what the agent believed to be the best option and the actual outcome. In both cases, this leads to an increase in *λ^∗^*, consistent with a possible change in contingencies.

The estimate is updated as a delta rule:

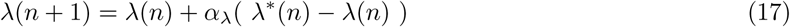

Meta-RL-incompatibility therefore has seven free parameters: *α*_0_, *a_λ_*, *α_λ_*, *a_ɛ_*, *α_ɛ_*, *β* and *κ*.

#### 2.5.4 Comparison models

To place the reinforcement-learning models in context, we compared them with three additional models that capture choice behavior without a value-based learning rule: the win-stay/lose-shift model defined above (Eq. 2, one free parameter *ɛ*), and the two models below.

##### Recursively formulated logistic regression (RFLR)

This model, adapted from [13], predicts choices from an exponentially filtered history of rewarded choices rather than from option values. Writing *c̄*(*n*) = +1 if option *L* was chosen on trial *n* and −1 otherwise, a recursive term *ϕ*(*n*) accumulates the reward-weighted choice history with time constant *τ* :

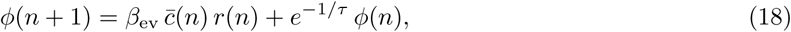

and the choice log-odds combine the last choice with this history:

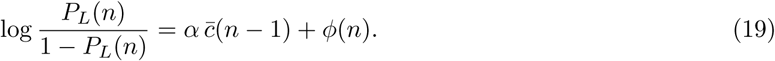

It has three free parameters: *α* (weight of the last choice), *β*_ev_ (reward sensitivity) and *τ* (history time constant).

##### State inference

This model, in the spirit of belief-based accounts [34], treats the task as inference over a hidden state indicating which option is currently best. The animal maintains a belief *b*(*n*) that option *L* is the better one. On each trial the observed outcome updates the belief by Bayes’ rule, using a reward-compatibility term (*c*_rew_ after a reward, *d*_unr_ after an omission) that measures how consistent the outcome is with the current belief, followed by a transition step with stay probability 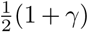 that allows the hidden state to reverse. Choices are drawn from a softmax on the belief with the same choice stickiness as the RL models. It has five free parameters: *c*_rew_, *d*_unr_, *γ*, *β* and *κ*.

### 2.6 Normative simulations

To derive the reward-maximizing parameters as a function of the environment (Fig. 2), we simulated the RL model with choice stickiness, using the coupled value update (Eq. 7), as an artificial agent over a plane of simulated environments. This stochasticity-volatility plane, used for all normative simulations, was defined on a 40 × 40 mesh. Stochasticity was varied through the reward magnitude *R* ∈ [0.80, 1.60], with the reward probability of the best option set to *P* = *Ē/R* so that the expected reward per option *Ē* = 0.8 was held constant across the plane (magnitude compensation, as in the task), the two options being mirror-symmetric (*p*_best_ = *Ē/R*, *p*_worst_ = (1− *Ē*)*/R*). Volatility was varied through the block length *L* ∈ [10, 210] trials between reversals. The stochasticity and volatility indices reported on the axes are *P* (1 − *P*)*R* and 1*/L*, respectively.

For each environment, agents performed 1000 trials with reversals every *L* trials, and the objective was the average reward per trial, estimated over 2000 independent agents. In the example environments (Fig. 2A–C), we swept the learning rate at a fixed inverse temperature and the inverse temperature at a fixed learning rate to display the reward landscape along each parameter. For the parameter maps, we performed a one-dimensional grid search in each cell: the optimal learning rate map (Fig. 2D) was obtained by sweeping *α* over [0.05, 1] (20 values) with the inverse temperature fixed (*β* = 2.5), and the optimal inverse temperature map (Fig. 2E) by sweeping *β* over [0.5, 20] (20 values) with the learning rate fixed (*α* = 0.4); the choice stickiness was fixed at *κ* = 2.0 throughout. Each map thus reports the reward-maximizing value of one parameter while the other is held constant, so that D and E describe agents adapting a single parameter. Maps were lightly smoothed for display (Gaussian filter, *σ* = 0.8) and overlaid with iso-performance and iso-parameter contours.

### 2.7 Parameters fitting

For each model, parameters were fitted with a standard maximization of likelihood [21, 22]. For a given experimental session and a given model, the likelihood is here defined as:

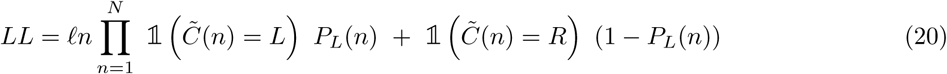

where 1 denotes the indicator function.

*C̃*(*n*) denotes the choice made by the animal at trial *n* in the given session and the variables of the model are updated based on the experimental reward. The first trial of each session was excluded from the likelihood, values being initialized at *V_L_*(0) = *V_R_*(0) = 0.5.

Minimization of the negative log-likelihood used the L-BFGS-B algorithm (*scipy.optimize.minimize*) within box constraints on each parameter. To limit sensitivity to local minima, each fit was repeated from 20 random initializations drawn uniformly within the bounds, and the solution with the lowest negative log-likelihood was retained. Learning rates and mixing probabilities were bounded to [0.01, 0.99], the inverse temperature to [0.1, 10], the choice stickiness *κ* to [0, 5], the incompatibility and stochasticity gains to [0.01, 2], and the RFLR history constant *τ* to [1, 100] trials.

The fitting was used in two ways. To characterize how the fitted learning rate and inverse temperature vary across the six environments (Fig. 4), the RL model with choice stickiness was fitted *per session*, i.e. separately in each of the six environments, so that *α*, *β* and *κ* could take a different value in each condition and their dependence on stochasticity and volatility be read directly from the fits. For model comparison, cross-validation, recovery and generative validation (Figs. 5–7), each model was instead fitted *per animal*, minimizing the negative log-likelihood summed over that animal’s six sessions, so that a single parameter set had to account for behavior across all six environments; the meta-RL models can express condition-dependence through their internal volatility and stochasticity estimates rather than through free per-condition parameters.

**Figure 4:**
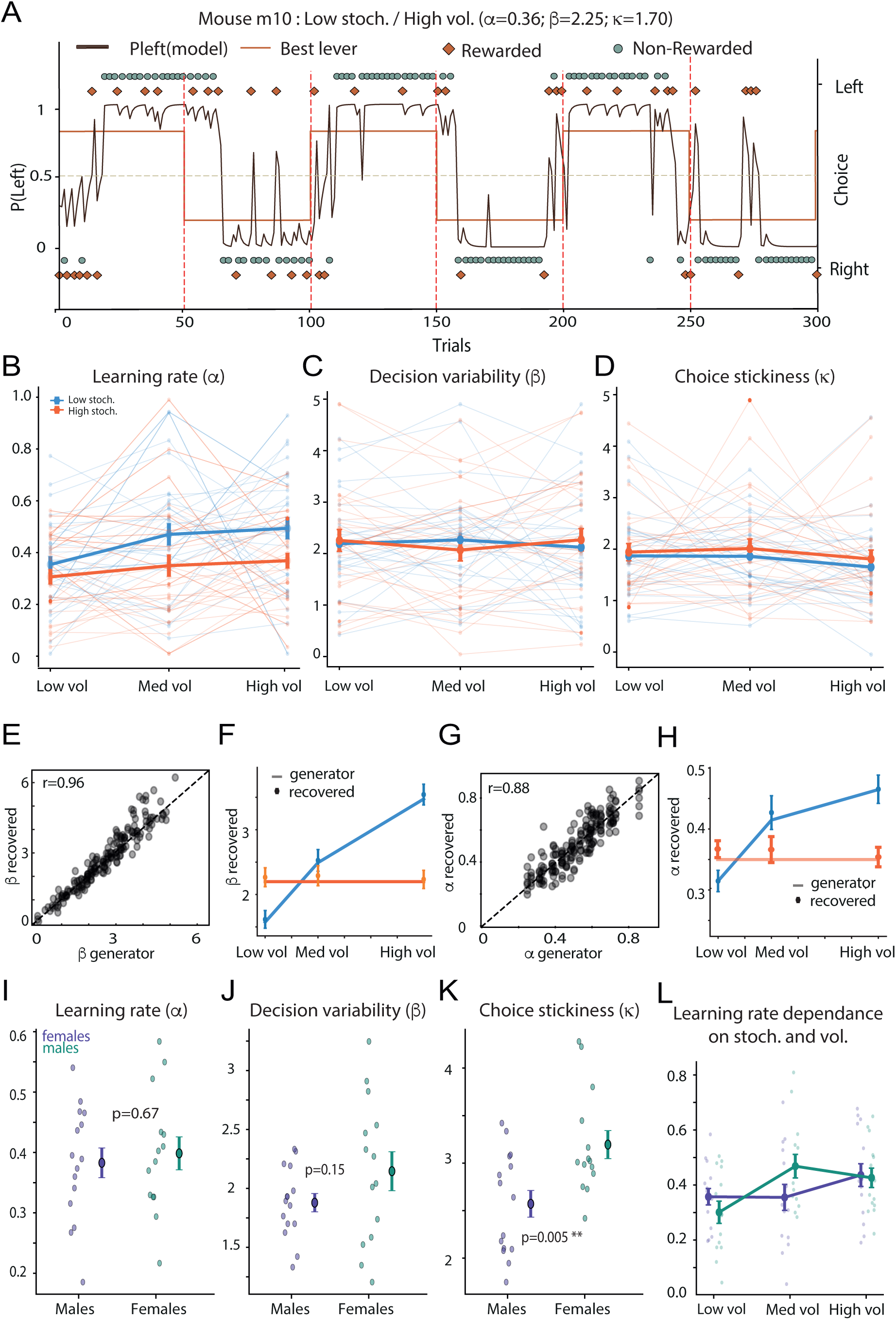
Fitted learning rates follow the normative predictions while inverse temperature is fixed. **(A)** Example model inversion: trial-by-trial probability of choosing the Left lever from the fitted RL model with choice stickiness (black line) overlaid on the animal’s choices (points), over several reversals in reward probability (orange line). **(B)** Fitted learning rate *α* as a function of volatility and stochasticity. *α* increases with volatility and decreases with stochasticity. **(C)** Fitted inverse temperature *β* across the same six environments. *β* is constant across conditions. **(D)** Fitted choice stickiness *κ* as a function of volatility and stochasticity. *κ* is constant across conditions. **(E)** *β* recovery: recovered vs generating *β* across all synthetic sessions (r = 0.96). **(F)** Recovered *β* along the volatility axis when agents were generated with a volatility-dependent *β* (blue) or a flat *β* (orange); the procedure recovers the generating pattern in each case. **(G)** *α* recovery: recovered vs generating *α* (r = 0.88). **(H)** Recovered *α* along the volatility axis at low and high stochasticity, tracking the generating values. **(I)** *α* did not differ between male and female mice (*F* (1, 27) = 0.18, *p* = 0.67). **(J)** *β* did not differ between male and female mice (*F* (1, 27) = 2.23, *p* = 0.15). **(K)** *κ* differed between male and female mice (*F* (1, 27) = 9.36, *p* = 0.005). **(L)** Learning rate *α* is sensitive to volatility in both male and female mice (*F* (2, 54) = 4.09, *p* = 0.02).

**Figure 5:**
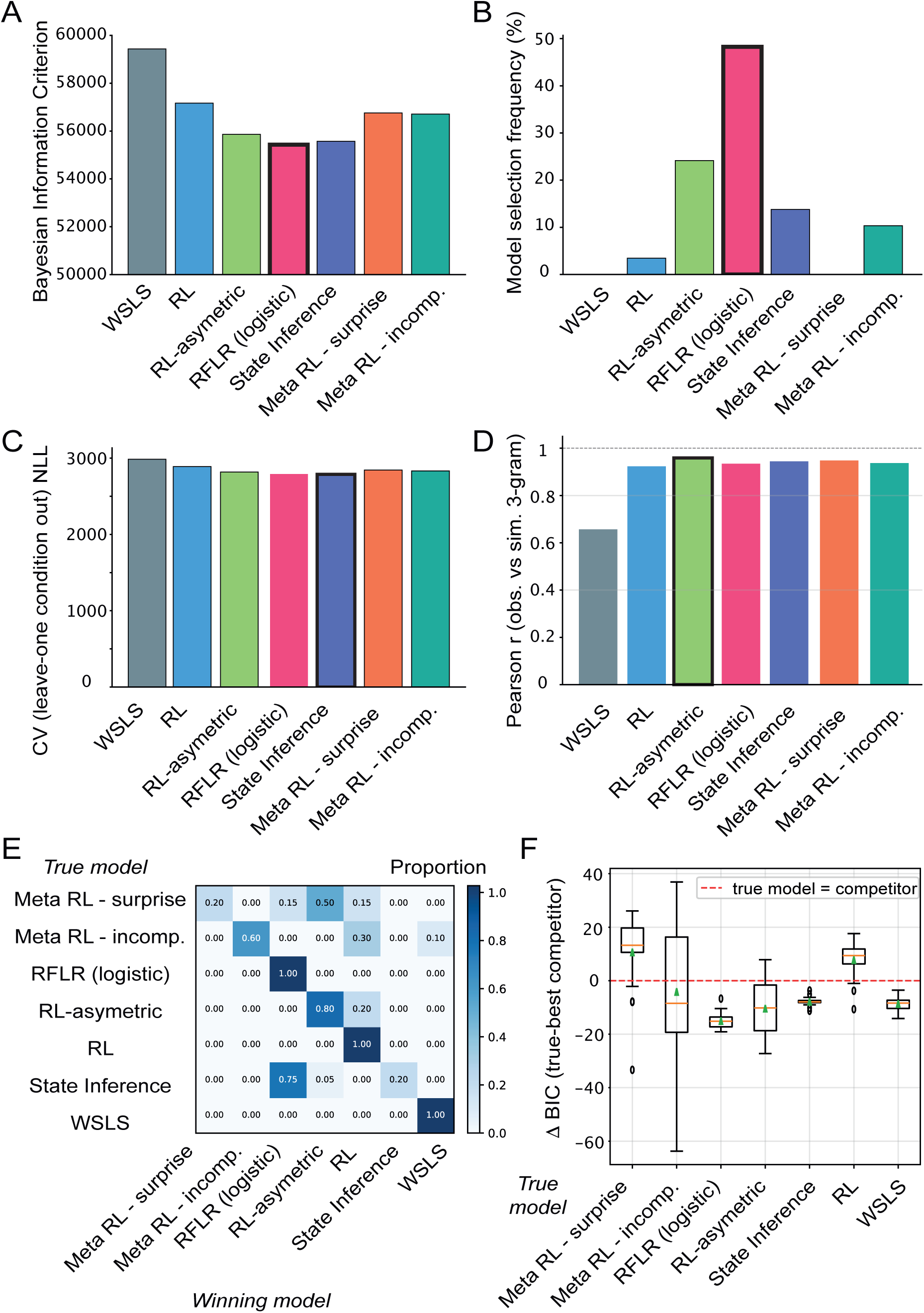
Standard fitting procedures fail to identify a single model capturing uncertainty-dependent learning. Panels A to D apply four selection criteria to the same fits, and the four disagree: group-level BIC favors RFLR (A), individual-level selection favors RFLR (B), leave-one-condition-out cross-validation favors state inference (C), and agreement with the empirical 3-grams favors RL-asymmetric (D). Model names follow Table 1. **(A)** Total Bayesian Information Criterion (BIC), i.e. model likelihood penalized by model complexity, was broadly similar across models and slightly favored RFLR (the favored model is indicated by a black outline). **(B)** RFLR is selected most frequently in individual fits. **(C)** Average negative log-likelihood (NLL) obtained by leave-one-condition-out cross-validation favors state inference. **(D)** Pearson correlation *r* between empirical 3-grams (Fig. 3H) and 3-grams simulated from each model favors RL-asymmetric. **(E)** Model recovery: each model, with parameters drawn from the animal fits, was simulated and refitted with the full model set. Cells give the proportion of winning models for each ground-truth model. **(F)** Difference in BIC in the recovery process between the true model and its best competitor. Negative values indicate favored recovery.

**Figure 6:**
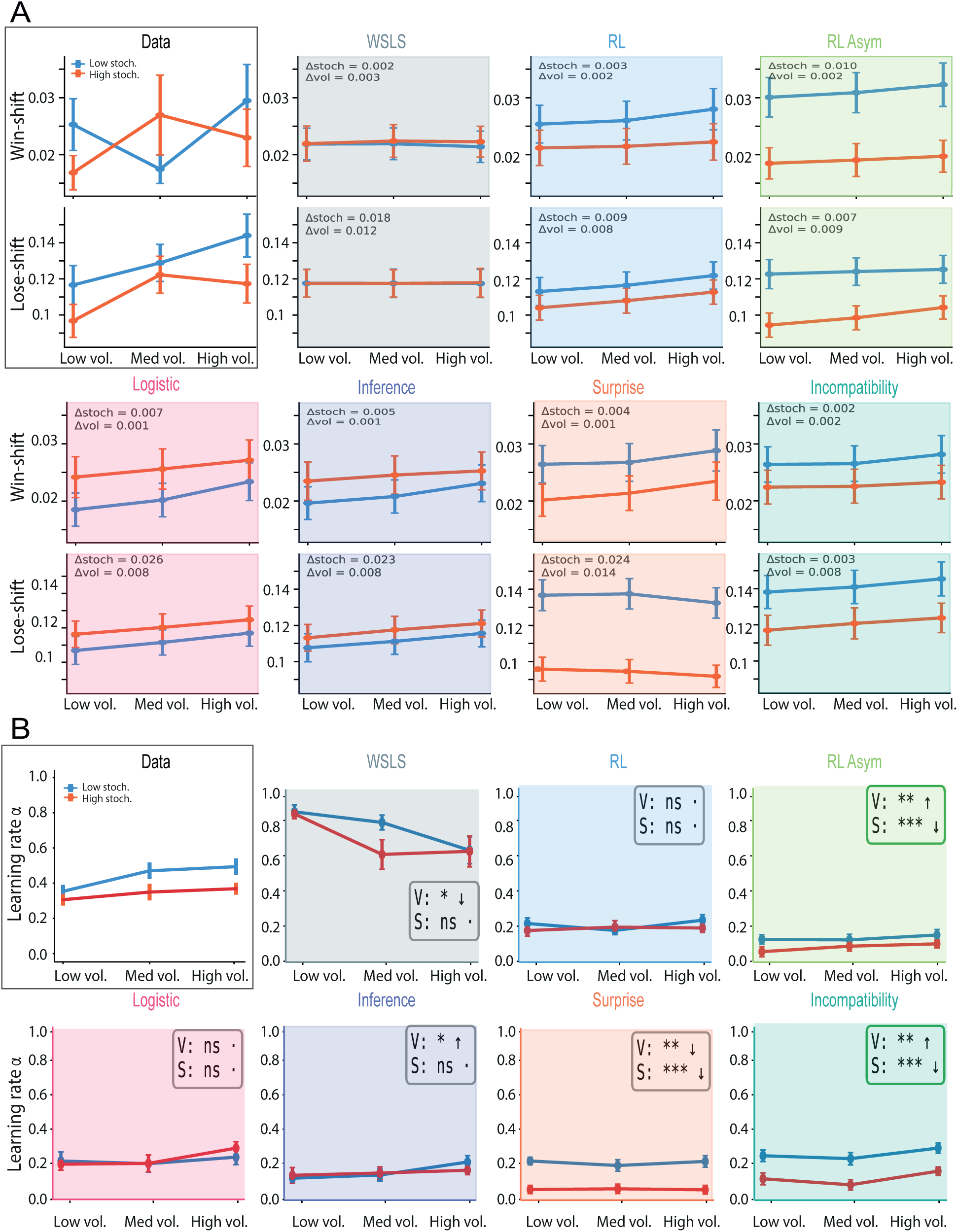
Only volatility-tracking models reproduce the adaptive learning signature of the data. (A) Win-shift (top row) and lose-shift (bottom row) as a function of volatility, at low (light) and high (dark) stochasticity, in the experimental data (leftmost panel, same as Fig. 3D,E) and in behavior simulated from each fitted model. Each model was simulated over the same six environments with the parameters fitted to each animal, 20 simulations per session. The data show a flat win-shift and a lose-shift that rises with volatility; the descriptive models (WSLS, RL, RFLR, state inference) reproduce the levels but not the gradient. (B) Effective learning rate *α*^, obtained by refitting the fixed-rate RL model to the simulated choices of each model, as a function of volatility at low and high stochasticity. Leftmost panel, the empirical *α*^ (same as Fig. 4B). Meta-RL-incompatibility reproduces both the increase with volatility and the decrease with stochasticity; meta-RL-surprise inverts the volatility dependence.

**Figure 7:**
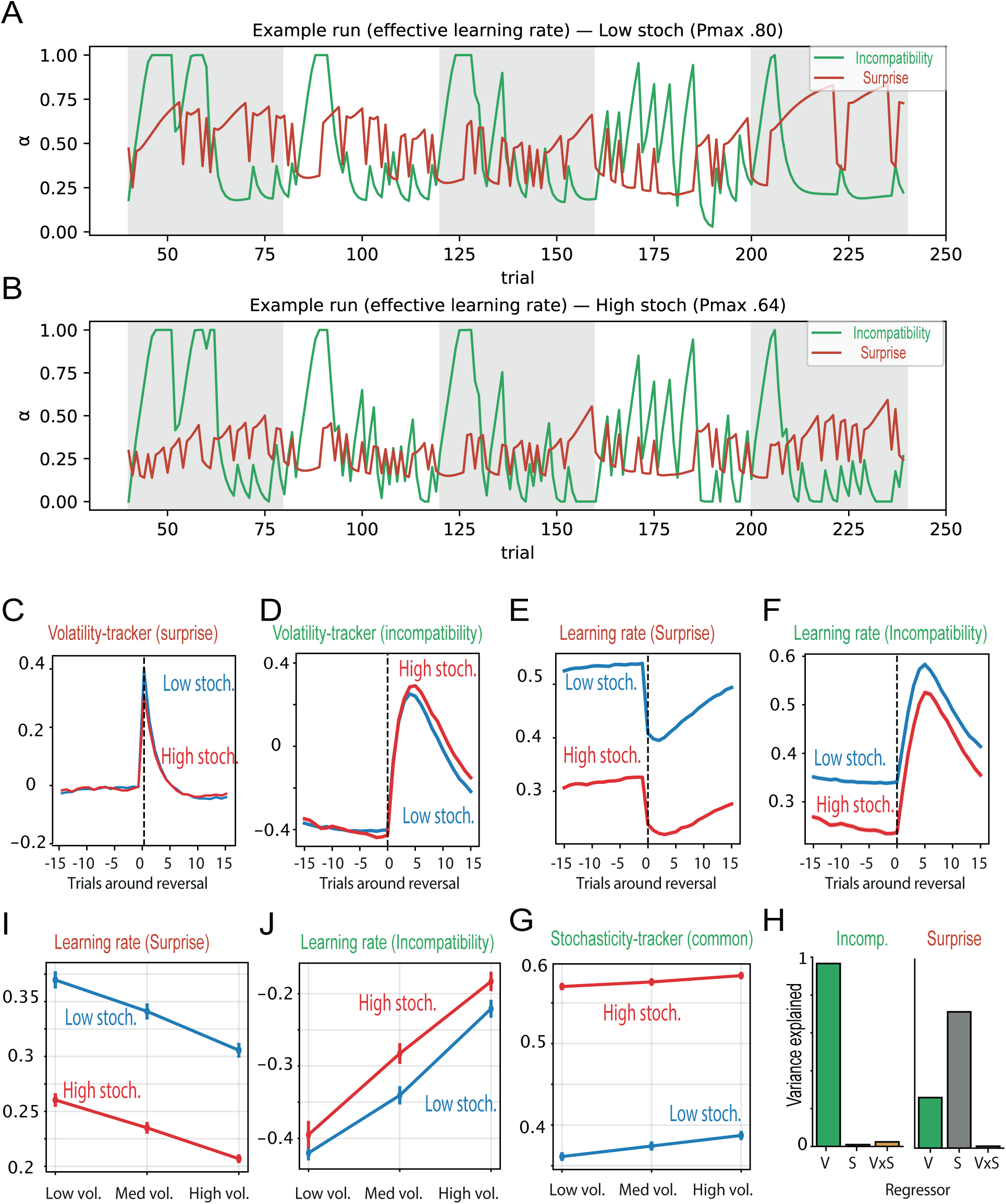
Volatility-tracking based on surprise is contaminated by stochasticity, contrary to the incompatibility heuristic. **(A,B)** Example simulations of the effective learning rate from the surprise and incompatibility meta-RL models under low (A) or high (B) stochasticity. **(C,D)** Reversal-centered volatility trackers based on surprise (unexpected uncertainty, C) and on incompatibility (D) both increase at reversal. **(E)** Reversal-centered learning rate driven by the surprise-based tracker: correct dependence on stochasticity but an inverted relation to volatility. **(F)** Reversal-centered learning rate driven by the incompatibility-based tracker: correct dependence on both stochasticity and volatility. **(G)** Stochasticity tracker (expected uncertainty), common to both models, depends mainly on stochasticity with little contamination from volatility. **(H)** Variance decomposition of the volatility trackers: volatility explains most of the variance of the incompatibility-based tracker, whereas the surprise-based tracker is dominated by stochasticity. **(I)** Session-averaged surprise-based learning rate decreases with volatility, opposite to the data. **(J)** Session-averaged incompatibility-based learning rate increases with volatility and decreases with stochasticity, matching the data.

**Table 1:** The seven models compared. *k* is the number of free parameters. The three families differ in whether the rule can change what it computes across the six environments: the non-adaptive rules cannot, the descriptive rules read the environment off the recent history, and the meta-learning rules maintain internal estimates of stochasticity and volatility and set the learning rate from them. All value-based models share the same softmax with choice stickiness *κ*, and all were fitted with a single parameter set per animal for the model comparison (Section 2.7).

| Model | Family | $k$ | Rule, and how its behavior can differ between environments |
| --- | --- | --- | --- |
| WSLS | non-adaptive | 1 | Repeat after a reward, switch after an omission, departing from the rule with probability $\epsilon$ (Eq. 2). Nothing in the rule responds to the environment. |
| RL | non-adaptive | 3 | Coupled Rescorla-Wagner update with a fixed learning rate $\alpha$, softmax with inverse temperature $\beta$ and choice stickiness $\kappa$ (Eqs. 4, 5, 7). Behavior differs between environments only because the outcomes do. |
| RL-asymmetric | non-adaptive | 4 | Same, with separate learning rates after rewards and omissions, $\alpha_+$ and $\alpha_-$ (Eq. 8). Its effective learning rate can shift with the proportion of rewarded trials, hence along the stochasticity axis, but no term responds to the rate of contingency change. |
| RFLR | descriptive | 3 | Choice log-odds from the last choice and an exponentially filtered history of rewarded choices, with weight $\alpha$, reward sensitivity $\beta_{ev}$ and time constant $\tau$ (Eqs. 18, 19). Reads the environment off the recent history instead of estimating it. |
| State inference | descriptive | 5 | Bayesian belief over which lever is currently better, updated by a reward-compatibility term ($c_{rew}$, $d_{unr}$) and a transition step ($\gamma$), then softmax with $\beta$ and $\kappa$. Requires the state space and the transition structure to be supplied. |
| Meta-RL-surprise | meta-learning | 7 | RL whose learning rate is $\alpha_0 \lambda(1 - \epsilon)$, with the stochasticity estimate $\epsilon$ and the volatility estimate $\lambda$ both read from the magnitude of prediction errors (Eqs. 9–13). |
| Meta-RL-incompatibility | meta-learning | 7 | RL whose learning rate is $\alpha_0 + a_\lambda \lambda - a_\epsilon \epsilon$, with $\epsilon$ read from the magnitude of prediction errors and $\lambda$ from the sign of the choice-outcome pairing (Eqs. 14–17). |

### 2.8 Model comparison

Models were compared with the Bayesian Information Criterion,

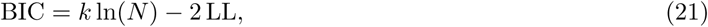

where *k* is the number of free parameters, *N* the total number of scored trials across the six sessions, and -LL the minimized negative log-likelihood. The seven models compared are listed in Table 1: WSLS (*k* = 1), RL with choice stickiness (*k* = 3), RL-asymmetric (*k* = 4), RFLR (*k* = 3), state inference (*k* = 5), meta-RL-surprise (*k* = 7) and meta-RL-incompatibility (*k* = 7). We report the group-level BIC (summed over animals), the frequency with which each model was selected at the individual level, and the pairwise ΔBIC against the best-fitting alternative.

#### Cross-validation

Because BIC penalizes complexity only through parameter count, we also assessed out-of-sample prediction with a leave-one-condition-out cross-validation. For each animal, the model was fitted on five of the six environments and its negative log-likelihood evaluated on the held-out environment, cycling over all six folds. The cross-validated NLL, summed over folds and animals, measures how well each model generalizes to an environment it was not fitted on.

#### Model recovery

To check that the models are distinguishable given our task and session lengths, we ran a model-recovery analysis. Each model was used in turn as a generator: synthetic animals were simulated with parameters drawn from the pool of fitted values, over the same six environments, and every model was then refitted to each synthetic dataset. The generating model was “recovered” when it obtained the lowest BIC. The resulting confusion matrix reports, for each generator, the proportion of synthetic datasets assigned to each model, and its diagonal quantifies identifiability.

### 2.9 Generative validation

Model comparison scores how well a model assigns likelihood to observed choices, but not whether the model, run forward, reproduces the qualitative behavioral signatures of the data. We therefore simulated each fitted model as a generative agent. Using each animal’s fitted parameters, we simulated behavior over the same six environments (matched block lengths, reward probabilities and magnitudes, and number of trials), sampling choices from the model’s own choice rule rather than replaying the animal’s choices. Each session was simulated 20 times and the resulting behavioral measures (win-shift, lose-shift, perseverative errors, post-reversal recovery) averaged, then compared with the empirical measures across the six environments (Fig. 6A). To recover an effective learning rate for each simulated agent, the simulated choices were refitted with the simple RL model, exactly as done for the real data, yielding the model-implied dependence of *α* on stochasticity and volatility (Fig. 6B).

As a further, choice-sequence-level test, we compared empirical and simulated *n*-gram statistics [13]. Each trial *n >* 0 was labelled by whether the animal repeated or switched its choice and whether that choice was rewarded, giving four symbols (rewarded stay, unrewarded stay, rewarded switch, unrewarded switch). For *n* = 2 and *n* = 3 we then computed the probability of switching on the next trial conditional on the preceding *n*-symbol history, both in the data and in sequences simulated from each model (one simulation per session, matching the empirical sample), and summarized the agreement by the Pearson correlation between observed and simulated conditional switch probabilities across histories and conditions (Fig. 5D).

### 2.10 Latent volatility and stochasticity tracking

To understand why the two meta-RL models differ in their volatility dependence (Fig. 7), we examined their latent trackers directly. For a given parameter set, we simulated each model over a 9 × 9 mesh of simulated environments spanning the experimental range of stochasticity and volatility (30 agents of 1500 trials per cell), and read out the steady-state value of each latent variable, discarding the first half of each session as burn-in. This was done for the volatility tracker *λ* and the stochasticity tracker *ɛ* of each model, using group-mean fitted parameters.

For each latent map we quantified how strongly the tracker depended on each environmental axis in two ways. First, we computed the conditional slope of the latent with respect to volatility at each fixed stochasticity level (linear fit on standardized volatility), which gives the sign and magnitude of the volatility dependence independently of stochasticity. Second, we partitioned the variance of the latent map across the simulated plane into a stochasticity main effect 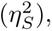 a volatility main effect 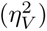 and their interaction 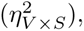 as fractions of the total sum of squares. A near-pure volatility tracker has 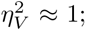 a tracker contaminated by stochasticity has a substantial 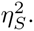 Reversal-centered trajectories of the effective learning rate (Fig. 7C-F) were obtained by aligning simulated trials on reversals and averaging within stochasticity levels, and the peri-reversal excursion was measured as the change in effective learning rate from baseline to its post-reversal peak.

### 2.11 Statistical analysis

The behavioral data collected across the different experimental environments were analyzed using two-way repeated-measures ANOVAs (RM-ANOVAs), with environmental stochasticity and volatility as within-subject factors. The parameters estimated through model fitting were analyzed in the same way. These analyses were performed in Python using the *SciPy* library (*scipy.stats* module).

Sex differences were assessed in a separate, exploratory set of analyses, for which the study was not specifically powered [36]. For these, sex was entered as a between-subject factor [43] alongside the within-subject factors (stochasticity, volatility) in mixed-design ANOVAs computed with the *pingouin* library. With 15 male and 14 female mice, between-subject effects have 1 and 27 degrees of freedom, and the three-level volatility factor has 2 and 54 degrees of freedom. We report these results as robustness checks rather than as confirmatory tests.

### 2.12 Data and code availability

The behavioral data and the analysis and modeling code supporting this study will be made available in a public repository upon publication.

## 3 Results

### 3.1 Compensating probability with magnitude separates stochasticity from task difficulty

Isolating stochasticity experimentally is harder than it appears, because a manipulation intended to vary it moves other important quantities along with it. Stochasticity is the variance of outcomes around a fixed reward expectation, and varying it should leave the value of an option unchanged. A study that varies stochasticity must therefore keep the total reward on offer constant, and the standard design does this by trading one option’s probability against the other (*p_R_* = 1 − *p_L_*). Along this path, while outcome variance changes as intended (Fig. 1A), the total value available is held fixed (Fig. 1B). The same path, however, cuts across the value difference between the two options, the quantity that sets how hard they are to tell apart (Fig. 1C). Here we define task difficulty as the difference in expected reward between options, Δ*E* = *E*(*R_left_*) − *E*(*R_right_*), the quantity that sets how discriminable they are. Variance and difficulty move together under this design (Fig. 1D), so a change in behavior across such conditions cannot be assigned to stochasticity rather than to difficulty. Compensating probability (*P*) with reward magnitude (*M* ) such that

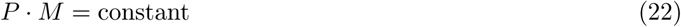

likewise varies outcome variance (Fig. 1E) while holding total value fixed (Fig. 1F). Value difference along this compensated path is held constant as well (Fig. 1G).

The two experimental conditions used in the rest of the manuscript thus sit on this compensated path: 80/20 with *M* = 1 and 64/16 with *M* = 1.25 both hold the expected reward of each option at 0.8 and 0.2, and their difference at 0.6, while the outcome variance of the better option rises from 0.16 to 0.36 (Fig. 1E–H, overlaid points; see also Fig. 3). The two options remain equally discriminable, so changes in outcome noise selectively alter variance (Fig. 1H).

Rodent studies of learning under uncertainty have varied reward probability without this compensation, so that outcome variance and option discriminability moved together [41, 13, 11], and have treated block length as a quantity drawn at random from block to block rather than as a factor [41, 11]. Hence, to our knowledge the design used here is the first in rodents in which outcome variance is varied at constant expected reward and constant value difference, and crossed factorially with volatility, which is what allows the effects reported below to be attributed to outcome variance itself.

### 3.2 Either the learning rate or the decision policy can be tuned to maximize reward

We then asked what an optimal reinforcement-learning agent *should* do in each of our environments. We simulated the RL model with choice stickiness (Eqs. 4, 5 and 7) and, for each environment, estimated the average reward per trial as a function of the model’s parameters, sweeping the learning rate at a fixed inverse temperature and the inverse temperature at a fixed learning rate (Fig. 2A, B, C). Each example environment yields a reward landscape whose peak defines the parameter value that maximizes reward, i.e. the *normative benchmark* against which behavior can be compared.

Repeating this over the full stochasticity-volatility plane gives, for each simulated environment, the reward-maximizing learning rate obtained while the inverse temperature is held at a fixed value, and the reward-maximizing inverse temperature obtained while the learning rate is held at a fixed value. The optimal learning rate *α*^opt^ rises with volatility and falls with stochasticity (Fig. 2D). When the environment is volatile, an unexpected outcome is more likely to signal that contingencies have genuinely changed, so the agent should weight it heavily and update quickly; when the environment is stochastic, the same outcome is more likely to be noise, so the agent should discount it and update slowly [8].

The optimal inverse temperature *β*^opt^ behaves similarly (Fig. 2E). It increases with volatility, and decreases with stochasticity. Yet the reward landscape is shallow along the *β* axis (Fig. 2A–C), so that a wide range of temperatures yields near-identical performance in any given environment. An agent that moves only its learning rate (inverse temperature fixed; Fig. 2D) and an agent that moves only its inverse temperature (learning rate fixed; Fig. 2E) both recover nearly all of the reward available across the plane: averaged over the plane, the agent that tunes only its learning rate at *β* = 2.5 leaves 0.094 reward per trial unclaimed, and the agent that tunes only its inverse temperature at *α* = 0.4 leaves 0.108, a difference of 0.014 reward per trial.

If animals follow a principle of reward maximization they need to fine-tune one of the two parameters, but the normative analysis does not prescribe which, so we next test which one mice actually move (Fig. 4).

### 3.3 Mice specifically adjust their response to losses across the stochasticity and volatility combinations

We trained mice in a binary reversal task (Fig. 3A)in which the two levers delivered intracranial self-stimulation (ICSS) of the medial forebrain bundle, because the reward is delivered directly, satiety does not accumulate over a session, motivation stays high and comparable across animals, sessions, and the thousands of trials needed to estimate choice statistics within each environment [28, 29, 30, 31].

On this paradigm we crossed our two sources of uncertainty in a 2 × 3 factorial design (Fig. 3B), with 29 mice contributing one session per environment and 144,647 trials in total. Stochasticity took two levels, set by the compensated probability pairs (80*/*20 and 64*/*16); volatility took three levels, set by the block length over which contingencies were held fixed before reversing (25, 50, and 100 presses). These six environments sample the region of the stochasticity-volatility plane simulated in Fig. 2, and let us ask whether behavior moves across them as the normative analysis predicts.

Success rate, the proportion of presses that were rewarded, was highest in stable, low-stochasticity environments and declined as either source of uncertainty grew (Fig. 3C), as the simulated agents do (Fig. 2A–C). Both factors contributed: success rate depended on stochasticity (*F* (1, 28) = 11.92, *p* = 0.0018) and, more strongly, on volatility (*F* (2, 56) = 78.66, *p <* 10*^−^*^9^; two-way repeated-measures ANOVA). Mice are thus sensitive to both manipulations, and in the predicted direction.

The two manipulations act on choice asymmetrically. Win-shift behavior, i.e. abandoning a lever that has just delivered reward, was flat across the six environments, depending neither on stochasticity (*F* (1, 28) = 0.21, *p* = 0.65) nor on volatility (*F* (2, 56) = 0.83, *p* = 0.44; Fig. 3D). Lose-shift behavior, by contrast, was strongly graded by the environment, the probability of switching after an unrewarded press rising with volatility (*F* (2, 56) = 4.61, *p* = 0.017) and decreasing with stochasticity (*F* (1, 28) = 7.63, *p* = 0.010; Fig. 3E). Mice therefore adapt how they respond to losses while leaving their response to wins essentially untouched.

The same conclusion emerges from the dynamics around a reversal. We counted the consecutive errors a mouse made immediately after contingencies changed, before it settled on the new best lever, a standard index of flexibility in reversal-learning tasks [27, 37] (Fig. 3F, G). These perseverative errors grew with stochasticity (*F* (1, 28) = 4.63, *p* = 0.040) : when outcomes are noisy, a run of omissions is weaker evidence that the world has changed, so recovery is slower. Perseverative errors also strongly depended on volatility (*F* (2, 56) = 7.61, *p* = 0.0026). Recovery curves thus separated across environments in the way a modulated learning rate, but not a fixed one, produces. The graded recovery seen here suggests that the effective learning rate itself changes with the environment. We decomposed switching by recent outcome history (Figure 3H), computing the probability of switching levers conditional on the preceding sequence of choices and rewards (n-gram analysis, following [13]). Across conditions, switch probability was structured by history rather than fixed: sequences ending in unrewarded presses on the current lever drove switching most strongly, and this structure deepened with volatility. The dependence is again asymmetric: the probability of switching after a win (win-shift) was low and insensitive to the environment, while switching after a loss (lose-shift) was strongly graded.

Finally, because block length was fixed within a session, mice could in principle have timed their switches to the block rather than to the outcomes. They did not. The probability of switching over the five presses preceding a reversal did not differ from its value during the stable part of the block (paired *t*(28) = −0.15, *p* = 0.88; Fig. 3I). The adaptation reported here is driven by feedback rather than by an internal count of presses.

We also checked whether male and female mice differed in reinforcement-learning properties [44] depending on the uncertainty of the environment. We found that their success rate was not different (Fig. 3J), yet male and female mice systematically differed in win-shift (Fig. 3K) and lose-shift (Fig. 3L), with female mice displaying less shift for all volatility schedules. Perseverative errors followed the same asymmetry: they were more numerous in female than in male mice (*F* (1, 27) = 6.48, *p* = 0.017) and increased with volatility in both sexes (*F* (2, 54) = 8.03, *p <* 0.001; Fig. 3M). Female mice thus shifted less after both wins and losses and took longer to leave the previously rewarded lever, a set of differences consistent with a single change in the tendency to repeat the last action rather than with a change in how outcomes are weighted.

Taken together, these observables provide a first, model-independent test of the normative predictions: performance falls with both kinds of uncertainty as predicted, and the adaptation is mainly carried by responses to losses. However, these model-free observables only establish that something is retuned with volatility, prompting us to identify whether the learning or the decision component carries this adjustment.

### 3.4 Mice adjust their learning rate, not their decision policy

We next asked whether the modulation of learning matches the normative predictions (Fig. 2). To answer this, we fitted the RL model with choice stickiness [13] (Eqs. 4, 5 and 7) to each animal’s choices, condition by condition, by maximizing the likelihood (Eq. 20). Fitting separately in each of the six environments lets the learning rate *α* and the inverse temperature *β* vary freely from one to the next. This fit is descriptive by construction: it measures how far the two parameters move, without saying what makes them move, which is the question taken up in the following sections. Inverting the model returned trial-by-trial choice probabilities that followed the animals’ actual lever sequences, including the dips and recoveries around reversals (Fig. 4A).

Read across the six environments, the fitted learning rate behaved as the normative analysis predicts (Fig. 4B). *α* increased with volatility : mice updated faster when contingencies turned over more often, and decreased with stochasticity, i.e. they updated more slowly when outcomes were noisier. Both dependencies match the *α*^opt^ map in sign (Fig. 2D): the learning rate is set jointly by the two statistics of the environment, in opposite directions.

By contrast, the fitted inverse temperature (*β*) was essentially constant across the six environments, moving with neither stochasticity nor volatility (as well as *κ* Fig. 4C,D). Recall that the reward landscape is relatively shallow along the *β* axis (Fig. 2A–C): over the range of temperatures the mice occupy, retuning *β* would buy almost no additional reward. An agent that holds its decision policy fixed and adapts only its learning rate therefore sits close to optimal, and that is the strategy the mice adopt.

The stability of *β* raises an identifiability concern : because the reward landscape is shallow along *β* (Fig. 2A–C), the likelihood is only weakly informative about this parameter. A flat *β* estimate could in principle reflect an estimator that cannot see *β* rather than an animal that holds it fixed. If the per-condition fitting procedure were blind to *β*, it would return a flat *β* whatever the animal did. Agents were thus simulated and refitted condition by condition exactly as the real data. Across the full range, *β* was recovered with high fidelity (r = 0.96; Fig. 4E). Decisively, the procedure recovered flat or increasing patterns of *β* (Fig. 4F). The learning rate passed the same test (Fig. 4G, H). This recovery strengthens the case that animals actually displayed a graded *α* and a flat *β*.

We also checked whether reward magnitude *M* could produce apparent changes in the fitted parameters, reflecting a change of reward units rather than a change in the underlying policy. However, writing the outcome as *r* = *Mb* with *b* ∈ {0, 1} shows that the value update (Eqs. 6 or 7) is homogeneous in *M*, so the learning rate, which multiplies a prediction error expressed in the same units as the value it corrects, is dimensionless and unchanged by the compensation, whereas the softmax argument *β*Δ*V* scales with *M*. An animal holding its policy fixed in normalized units would therefore return the same *α* in the two stochasticity conditions and a *β* lowered by a factor 1*/*1.25 at high stochasticity. Neither prediction is met: *α* falls with stochasticity (Fig. 4B), and *β* does not show the predicted displacement along that axis (Fig. 4C). Hence, the dependence of the learning rate on the environment is not confounded by the reward magnitude.

Finally, we asked whether these parameters differed between male and female mice. The two adaptive parameters did not differ by sex. Neither the learning rate *α* (F(1, 27) = 0.18, p = 0.67; Fig. 4I) nor the inverse temperature *β* (F(1, 27) = 2.23, p = 0.15; Fig. 4J) separated the sexes. Choice stickiness *κ* did: females carried a higher choice stickiness than males (F(1, 27) = 9.36, p = 0.005; Fig. 4K). The volatility dependence of the learning rate held in males and females alike (F(2, 54) = 4.09, p = 0.02; Fig. 4L), with no interaction between sex and the adaptive tuning of *α*. The sex difference is thus confined to a parameter that governs how readily a mouse repeats its last action, not to the parameters through which it adapts to uncertainty.

Overall, the learning rate follows the *α*^opt^ map in both of its dependencies, which sets up the question of how mice move their learning rate this way.

### 3.5 Standard model fitting cannot identify the rule that generates the learning-rate adaptation

Having established that mice adapt their learning rate to volatility and stochasticity (Fig. 4), we asked which learning rule generates this adaptation. We compared seven models spanning three families (Methods, Section 2.5): rules with no adaptive machinery (win-stay/lose-shift, fixed-rate RL, asymmetric RL), descriptive rules that read the next choice off recent history (RFLR, state inference), and meta-learning rules that compute a trial-by-trial learning rate from internal estimates of stochasticity and volatility (meta-RL-surprise, meta-RL-incompatibility). Table 1 summarizes the seven. The two meta-learning rules share an architecture and differ only in how volatility is estimated, from the magnitude of prediction errors [41] or from the sign of successive choice-outcome pairs [33]. A full Bayesian joint estimator was not included, since by definition it recovers the reward-maximizing learning rate from the task structure it is handed [8].

All seven models were fitted per animal, a single parameter set accounting for the six environments (Methods), so that no model could carry the cross-environment structure in condition-specific parameters. On this common footing RFLR obtained the best group-level Bayesian Information Criterion, with the two meta-RL models within 100 points of it on a group-level score of 55,000 (Fig. 5A), and was selected in 14 of 29 animals (Fig. 5B). The ranking did not survive a change of criterion. Leave-one-condition-out cross-validation returned state inference (Fig. 5C), and the *n*-gram structure of the choices was reproduced with a Pearson *r* above 0.9 by every model except WSLS (Fig. 5D). No two criteria agreed on which model was best.

Whether such a disagreement reflects a genuine tie between the models or a limit of the selection procedure is a question about identifiability, which we addressed with a model-recovery analysis run for that purpose: each model was simulated with the fitted parameters and the full set was refitted to the synthetic data. The descriptive and non-adaptive rules were recovered essentially perfectly, RFLR, RL and WSLS each in 100% of simulations (Fig. 5E). The adaptive rules were not. Data generated by meta-RL-incompatibility were recovered as that model in 60% of simulations and assigned to RL in 30%; data generated by meta-RL-surprise were recovered in 20% of simulations and assigned to RL-asymmetric in 50%. The ΔBIC against the best competitor was accordingly negative for every descriptive model and positive for both meta-RL models (Fig. 5F). The confusion is thus not symmetric: it is confined to the adaptive models and directed toward the descriptive ones, which is the pattern expected if the two families are not on an equal footing under this criterion.

The direction of that confusion follows from what a likelihood measures. The likelihood, and therefore the BIC, is a one-step-ahead quantity: on every trial the model is handed the animal’s own sequence of past choices and outcomes, and is scored only on the choice that comes next. That history already carries the imprint of the environment the animal was in, so a model that filters it can reproduce the differences between environments without ever computing stochasticity or volatility. A model that does compute them online must additionally get their dynamics right, and pays for the attempt in free parameters. Deciding which rule generates the adaptation therefore requires running each model forward and asking whether the adaptation reappears [23], which we do next.

### 3.6 Only a modulated learning rate generates the behavioral sensitivity to losses

The behavioral signature of learning-rate modulation, a win-shift that is flat across the six environments and a lose-shift that grades with volatility (Fig. 3D,E), is a model-independent observation. To test which learning rule generates it, we simulated each fitted model over the same six environments. Only the volatility-tracking rules reproduced both the asymmetry between wins and losses and its ordering across stochasticity levels; the descriptive rules matched the marginal levels of win-shift and lose-shift but left them flat across environments (Fig. 6A).

We then refitted each model’s synthetic choices with the simple RL model, exactly as done for the real data, and asked whether the resulting effective learning rate *α*^ carries the dependence observed in the animals. *α*^ is the learning rate that a fixed-rate RL model returns when fitted to a set of choices, that is the same quantity measured in the mice in Fig. 4B, and it can be read from any generative model whether or not that model has a learning rate of its own. Only meta-RL-incompatibility recovered both effects in the correct direction and with a comparable magnitude (volatility, *p <* 0.01; stochasticity, *p <* 0.001; Fig. 6B). RL-asymmetric moved *α*^ in the right direction but too weakly, which is expected from its structure: it has no term that responds to the rate of contingency change, and its effective rate can shift only through the proportion of rewarded trials, that is along the stochasticity axis. Meta-RL-surprise failed qualitatively, its effective learning rate decreasing with volatility, opposite to the normative and observed pattern. The remaining models showed no reliable dependence on either axis.

Overall, only meta-RL-incompatibility reproduced the sign and the magnitude of the effects on both the model-independent and the model-dependent measures, while meta-RL-surprise produced the opposite pattern. We next examine the latent variables that produce this separation.

### 3.7 An incompatibility-based heuristic is volatility-specific, unlike a surprise-based signal

The refit showed that meta-RL-surprise carries the wrong dependence of the learning rate on volatility (Fig. 6B). Both meta-learning rules raise the learning rate with an internal volatility estimate, so the failure must lie in what that estimate measures. We therefore examined the latent variables directly, simulating both models over the stochasticity-volatility plane across the experimental range and reading out the steady-state value of each tracker in each cell.

A volatility tracker faces a structural difficulty [8]. The only signal available to it is the stream of outcomes, and stochasticity and volatility both make that stream less predictable. Distinguishing them requires a quantity that responds to how often the sign of the evidence turns over, and not to how large the individual surprises are. Yet meta-RL-surprise reads both stochasticity and volatility from the magnitude of prediction errors, and tries to separate “expected” (stochasticity) from “unexpected” (volatility) uncertainty on that single signal. Meta-RL-incompatibility instead reads volatility from the sign of the choice-outcome pairing, whether repeating a choice was then unrewarded or switching was then rewarded.

The consequences are visible around a reversal (Fig. 7A,B). Both trackers rise when contingencies change (Fig. 7C,D), but they drive the learning rate in opposite ways. We measured the peri-reversal excursion as the change in effective learning rate from its pre-reversal baseline to its post-reversal peak. Under the incompatibility tracker this excursion is Δ*α*^ = +0.27 and is nearly the same at both stochasticity levels (Fig. 7F). Under the surprise tracker it is Δ*α*^ = −0.09, the learning rate falling rather than rising at the moment the environment changes, and separating by stochasticity instead (Fig. 7E).

Partitioning the variance of each tracker across the simulated plane, into a stochasticity main effect, a volatility main effect and their interaction, confirms the dissociation at the level of the environment statistics rather than of individual animals. The incompatibility tracker *λ* is a near-pure index of volatility 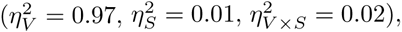 whereas the variance of the surprise tracker is carried mostly by stochasticity (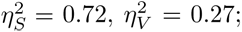 Fig. 7H). The stochasticity tracker, common to both models, separates stochasticity with little contamination by volatility (Fig. 7G). Session-averaged, the surprise-based *α*^ therefore decreases with volatility (Fig. 7I) while the incompatibility-based *α*^ increases with volatility and decreases with stochasticity, matching the data (Fig. 7J).

## 4 Discussion

### 4.1 The learning rate carries the adaptation to different forms of uncertainty

Across the six environments, mice raised their learning rate with volatility and lowered it with stochasticity, in both directions predicted by the normative analysis, while their inverse temperature stayed fixed [13, 41, 8].

Increases in learning rate under volatility are reported consistently, whereas the matching decrease under stochasticity has appeared mainly when the level of noise was signaled explicitly during the task [42] or when noise and volatility produced outcome ranges different enough to be told apart [45]. When they were made comparable, human participants adapted to volatility alone, and a Bayesian observer analysis indicated that they were responding to a coarse estimate of noise and attributing it to volatility [24, 46]. The same null result was obtained where the stochastic condition had been calibrated by moving the generative means of the two options together until difficulty matched the volatile condition, so that outcome variance and option discriminability changed at once [47]. The rodent and primate literature bear on the same question but does not separate the two statistics [11]. In both cases a reduction of learning rate under uncertainty is actually compatible with a response to task difficulty. By contrast, the magnitude compensation we introduce removes that reading. Our mice show a joint stochasticity-volatility effect, having received no description of the task structure (contrary to human studies), and in a condition where an increase in outcome variance cannot be read as the task becoming harder.

The stability of the inverse temperature is normatively grounded, since the reward landscape along *β* is relatively shallow (Fig. 2A–C), so an animal that leaves *β* where it was gives up almost no reward. Yet it is not the only solution the task would admit: an agent that had held its learning rate fixed while moving its decision policy would have performed comparably (Fig. 2D,E). Accordingly, adjustment of decision policy is a documented route to adaptation in its own right, with meta-learning models that regulate the exploration-exploitation balance by tuning the softmax temperature online from reward history or prediction error [17, 48, 49], and medial prefrontal and cingulate accounts that cast these regions as meta-learners adjusting reinforcement-learning meta-parameters from ongoing performance [50, 51, 52, 53]. An invariance of the same kind appears in continuous-time foraging, where the variance of the estimation error grew with the variance of reward timing while the variance of patch residence times did not [54]. Which parameter moves may therefore depend on the demands the task places on exploration, which are limited when the state space is small and the option set fixed [55, 56] like in our setup.

The sex comparison supports the same conclusion from an unplanned direction. Male and female mice differed in choice stickiness *κ* and not in *α* or *β*, and the volatility dependence of the learning rate held in both (Fig. 4I–L). The difference in the raw shift behavior is consistent with this assignment: female mice shifted less after both wins and losses, a joint displacement of the two shift types in the same direction, which is what a parameter governing choice repetition produces [39]. A difference in learning rate would grade the loss-driven shift while leaving the win-driven shift largely untouched, as volatility does across the six environments (Fig. 3D,E). Female mice have been reported to learn faster than males in a restless bandit task [44]; in our design the difference fell on choice repetition instead, consistent with sex acting on value-guided behavior through several partly separable computations [43]. Nevertheless, these analyses were exploratory and not specifically powered to detect these effects [36].

### 4.2 Heuristic, descriptive and normative accounts answer different questions

Normative, descriptive and heuristic accounts answer different questions, and their merits are not commensurable [57, 58]. A normative analysis fixes which effects should exist, setting the sign of the dependence of the learning rate on each source of uncertainty from first principles about what a surprise implies [8, 5, 59]. A descriptive model summarizes how behavior depends on recent history. Neither states how a biological agent, which never observes stochasticity or volatility directly, could arrive at those quantities from experience. Our contribution thus belongs to the third class. The heuristic commits to specific signals and update rules, unsigned prediction errors for stochasticity and a choice-outcome incompatibility term for volatility, and recovers the normative dependence of the learning rate on both statistics without representing a hidden state or its transition probabilities.

The heuristic model does not win on penalized likelihood, and the reason bears on how models are compared whenever behavior is measured across several environments. RFLR reached the lower BIC, and it did so with a single parameter set for the six environments [13]. The advantage does not come from extra flexibility but from what a likelihood measures. Every choice is scored conditional on the animal’s own history, in which the environment has already left its mark, so a model that filters that history reproduces the cross-environment differences without generating them. Penalized fit quality ranks models on how well they continue an observed sequence, not on whether they produce the structure of the experiment when run forward. Separating the two requires simulating each candidate and asking whether the effect of interest reappears [23, 22]. In our set, only meta-RL-incompatibility passed that test (Fig. 6B).

Yet, complexity is not by itself an argument in the other direction either. A hierarchical Bayesian account was recently favored for mice foraging under variable reward timing [54], but that estimator was supplied with the generative process, including the form of the reward distribution. The normative joint estimator likewise recovers the reward-maximizing learning rate from the task structure it is given, by construction [8, 60].

Hence, finding the proper low-complexity rule matters, since the isolation of volatility from stochasticity is a property of this heuristic and not of heuristics in general. Surprise is not the problem [61]: the normative estimator uses the same prediction-error stream, weighted by an online stochasticity estimate. What fails is the unweighted implementation, in which the volatility term is updated only after errors, so that how often it is updated depends on how often errors occur, and error frequency is set by stochasticity [41]. By contrast, the incompatibility-based volatility tracker is a near-pure index of volatility (Fig. 7H), which is what allows the reward-magnitude compensation to pull the two apart in the model as well as in the task.

### 4.3 The adaptive computation constrains the substrate only weakly

The heuristic isolates the computations without committing to their substrate, and the two signals it relies on map onto partly separable neuromodulatory and cortical systems [4, 6, 1, 62]. Adjusting the learning rate from an internal estimate of volatility has been tied to orbitofrontal noradrenaline, where release tracks trial-by-trial volatility and disrupting the noradrenergic input to orbitofrontal cortex reproduces the predicted deficit in learning-rate control [33, 63]. Consistent with a role for this region [64], orbitofrontal but not secondary motor cortex encodes choice and outcome more strongly as reward schedules become less certain, and its activity is predicted by win-stay and lose-shift behavior [65]. Whether the stochasticity term, read here from unsigned prediction errors, is carried by a distinct system remains open [66, 67]. The normative account predicts that it should be, since the two estimates pull the learning rate in opposite directions and are most useful when computed separately [8, 4].

The separation between a volatility signal and a stochasticity signal is a statement about the computation and does not require two dedicated systems. A single population of synapses could produce the same behavior, with slow metaplasticity [59, 6]. A synapse driven repeatedly in the same direction becomes progressively harder to move, whereas one recently pushed in both directions stays labile. Volatility or stochasticity-related neuromodulation and metaplastic, value-encoding synapses make the same global prediction on adaptive learning rate. They diverge when a single input pathway is manipulated in isolation: a two-signal architecture predicts that removing the pathway carrying volatility abolishes learning-rate adaptation while leaving value learning intact, whereas a purely synaptic implementation predicts no such dissociation [6, 68].

### 4.4 What the design leaves open

Three limits bound our conclusions. The first concerns the interpretation of the independence of *β* to volatility and stochasticity. An inverse temperature absorbs every source of variability in choice, and there are at least two. One is exploration proper, a policy that samples the lower-valued option on purpose. The other is imprecision in the value updates themselves, which makes choices variable without any policy being tuned [69]. A single inverse temperature cannot separate them, and the two push it in opposite directions: more exploration lowers *β*, less imprecision raises the apparent *β*. In humans, exploration rises and learning imprecision falls with volatility, and the two are regulated independently [47], so a model carrying a single inverse temperature registers their sum and returns a flat *β* when they offset. The recovery analysis rules out a failure of the estimator, since a graded *β* was recovered whenever one was present in the simulated data (Fig.4 E,F), but no fit of this form can distinguish a policy that is genuinely fixed from two sources of variability that cancel. What this would change is the interpretation rather than the result: the adaptation would still be carried by the learning rate, but the constancy of the policy would be apparent rather than real. The same caution applies to the constancy of win-shift, which sits near floor across the six environments and therefore constrains the mechanism only weakly; the informative observable is the gradient of lose-shift.

Second, the task presents a single pair of levers whose contingencies reverse, and the reward is intracranial self-stimulation. This holds motivation constant across the thousands of trials the design requires, and it leaves the animal with the smallest possible problem to solve. Adjusting the learning rate is sufficient in that regime. With more options, or with structure the animal must infer at several levels, exploration has to be aimed at particular options instead of being raised or lowered globally, and the stability of the inverse temperature we report need not hold [55, 70, 71].

Third, the magnitude compensation assumes that twice as many stimulation pulses are worth twice as much to the animal. Raising magnitude as probability falls keeps expected reward equal across environments only if subjective value grows linearly with pulse count. Over the narrow range used here (*M* = 1.00 against *M* = 1.25) this is a reasonable approximation, and the press counts did not differ between the two magnitudes (Section 3.3). The compensation was also tested, as a purely metrological account of the two conditions predicts a fixed displacement of *β* and none of *α*, which is the opposite of the observed pattern. Nevertheless, over a wider range subjective value saturates [28], and the compensated design would reintroduce the difference in discriminability it is built to remove.

By manipulating the two dimensions of uncertainty independently, this framework also provides a handle on disordered adaptation, where a deficit in estimating one dimension can present as an abnormality in the other [8, 72]. The capacity to match the learning rate to the statistics of the environment is a core component of behavioural flexibility, whose disruption is a transdiagnostic feature of psychiatric and neurodevelopmental conditions [73, 11]. Computational work in humans [74] has linked this disruption specifically to the estimation of volatility, with autistic adults overestimating how quickly the world changes [75] and noradrenergic blockade slowing the adjustment of learning to volatility [76]. Isolating stochasticity from volatility, and identifying the heuristics that track each of them in animals, offers a tractable entry point into how adaptive learning breaks down in these conditions.

## 5 Acknowledgement

We thank the IGF animal facility and the “Physiopathology of neuronal plasticity” lab for scientific discussions.

## 6 Author Contribution

Conceptualization : JN, Data curation : CC, JN, Funding acquisition : JP, JN, Investigation : CC, JC, ED, EM, JN, Methodology : CC, JC, JN, Project administration : JN, Resources : JP, JN, Software : CC, JC, JN, Supervision : JN, Validation : CC, JN, Visualization : CC, JC, Writing — original draft : CC, JN, and Writing — review & editing : CC, JC, ED, JP, JN.

## 7 Ethics

The study was approved by the CEEA local and national committees (APAFIS:2209-50949).

The authors declare no competing interest.

## References

[1] Ilya E. Monosov. “How Outcome Uncertainty Mediates Attention, Learning, and Decision-Making”. Trends in Neurosciences. doi: 10.1016/j.tins.2020.06.009.

[2] Peter Dayan and Nathaniel D. Daw. “Decision theory, reinforcement learning, and the brain”. Cognitive, Affective, & Behavioral Neuroscience. doi: 10.3758/CABN.8.4.429.

[3] Richard S Sutton and Andrew G Barto. “Reinforcement Learning: An Introduction”.

[4] Angela J. Yu and Peter Dayan. “Uncertainty, Neuromodulation, and Attention”. Neuron. doi: 10.1016/j.neuron.2005.04.026.

[5] Timothy E J Behrens et al. “Learning the value of information in an uncertain world”. Nature Neuroscience. doi: 10.1038/nn1954.

[6] Alireza Soltani and Alicia Izquierdo. “Adaptive learning under expected and unexpected uncertainty”. Nature Reviews Neuroscience. doi: 10.1038/s41583-019-0180-y.

[7] RA Rescorla and Allan Wagner. “A theory of Pavlovian conditioning: Variations in the effectiveness of reinforcement and nonreinforcement”. Classical Conditioning II: Current Research and Theory. Jan. 1972.

[8] Payam Piray and Nathaniel D. Daw. “A model for learning based on the joint estimation of stochasticity and volatility”. Nature Communications. doi: 10.1038/s41467-021-26731-9.

[9] Matthew R. Nassar et al. “An Approximately Bayesian Delta-Rule Model Explains the Dynamics of Belief Updating in a Changing Environment”. The Journal of Neuroscience. doi: 10.1523/jneurosci.0822-10.2010.

[10] Christoph Mathys. “A Bayesian foundation for individual learning under uncertainty”. Frontiers in Human Neuroscience. doi: 10.3389/fnhum.2011.00039.

[11] Jae Hyung Woo et al. “Mechanisms of adjustments to different types of uncertainty in the reward environment across mice and monkeys”. Cognitive, Affective, & Behavioral Neuroscience. doi: 10.3758/s13415-023-01089-1.

[12] Brian Lau and Paul W. Glimcher. “DYNAMIC RESPONSE-BY-RESPONSE MODELS OF MATCHING BEHAVIOR IN RHESUS MONKEYS”. Journal of the Experimental Analysis of Behavior. doi: 10.1901/jeab.2005.110-04.

[13] Celia C. Beron et al. “Mice exhibit stochastic and efficient action switching during probabilistic decision making”. Proceedings of the National Academy of Sciences. doi: 10.1073/pnas.2113961119.

[14] The International Brain Laboratory et al. “Standardized and reproducible measurement of decision-making in mice”. eLife. Ed. by Naoshige Uchida and Michael J Frank. doi: 10.7554/eLife.63711.

[15] Aaron C. Courville, Nathaniel D. Daw, and David S. Touretzky. “Bayesian theories of conditioning in a changing world”. Trends in Cognitive Sciences. doi: 10.1016/j.tics.2006.05.004.

[16] Robert C. Wilson, Matthew R. Nassar, and Joshua I. Gold. “Bayesian Online Learning of the Hazard Rate in Change-Point Problems”. Neural Computation. doi: 10.1162/NECO_a_00007.

[17] Kenji Doya. “Metalearning and neuromodulation”. Neural Networks. doi: 10.1016/S0893-6080(02)00044-8.

[18] Ryoma Hattori et al. “Meta-reinforcement learning via orbitofrontal cortex”. Nature Neuroscience.

[19] Payam Piray and Nathaniel D. Daw. “A simple model for learning in volatile environments”. PLOS Computational Biology. doi: 10.1371/journal.pcbi.1007963.

[20] Jennifer L Cook et al. “Catecholaminergic modulation of meta-learning”. eLife. doi: 10.7554/eLife.51439.

[21] Nathaniel D Daw. “Trial-by-trial data analysis using computational models”.

[22] Robert C Wilson and Anne GE Collins. “Ten simple rules for the computational modeling of behavioral data”. eLife. Ed. by Timothy E Behrens. doi: 10.7554/eLife.49547.

[23] Stefano Palminteri, Valentin Wyart, and Etienne Koechlin. “The Importance of Falsification in Computational Cognitive Modeling”. Trends in Cognitive Sciences. doi: 10.1016/j.tics.2017.03.011.

[24] Erdem Pulcu and Michael Browning. “Humans adapt rationally to approximate estimates of uncertainty”. Elife.

[25] Xiaotong Fang and Payam Piray. “Inferring the causes of noise from binary outcomes: A normative theory of learning under uncertainty”. bioRxiv.

[26] Roshan Cools et al. “Defining the Neural Mechanisms of Probabilistic Reversal Learning Using Event-Related Functional Magnetic Resonance Imaging”. Journal of Neuroscience. doi: 10.1523/JNEUROSCI.22-11-04563.2002.

[27] A. Izquierdo et al. “The neural basis of reversal learning: An updated perspective”. Neuroscience. doi: 10.1016/j.neuroscience.2016.03.021.

[28] William A. Carlezon and Elena H. Chartoff. “Intracranial self-stimulation (ICSS) in rodents to study the neurobiology of motivation”. Nature Protocols. doi: 10.1038/nprot.2007.441.

[29] Jérémie Naudé et al. “Nicotinic receptors in the ventral tegmental area promote uncertainty-seeking”. Nature neuroscience.

[30] Marwen Belkaid et al. “Mice adaptively generate choice variability in a deterministic task”. Communications Biology. doi: 10.1038/s42003-020-0759-x.

[31] Elise Bousseyrol et al. “Dopaminergic and prefrontal dynamics co-determine mouse decisions in a spatial gambling task”. Cell Reports.

[32] Nathaniel D. Daw et al. “Cortical substrates for exploratory decisions in humans”. Nature. doi: 10.1038/nature04766.

[33] Hadrien Plat et al. “Orbitofrontal noradrenaline supports adaptive learning-rate adjustment in probabilistic reversal learning”. Proceedings of the National Academy of Sciences.

[34] Karyna Mishchanchuk et al. “Hidden state inference requires abstract contextual representations in the ventral hippocampus”. Science. doi: 10.1126/science.adq5874.

[35] George Paxinos and Keth B J Franklin. “The Mouse Brain in Stereotaxic Coordinates”.

[36] Payam Piray. “Addressing low statistical power in computational modelling studies in psychology and neuroscience”. Nature Human Behaviour.

[37] Carl Harris et al. “Unique features of stimulus-based probabilistic reversal learning”. Behavioral Neuroscience. doi: 10.1037/bne0000474.

[38] Makoto Ito and Kenji Doya. “Validation of decision-making models and analysis of decision variables in the rat basal ganglia”. The Journal of neuroscience.

[39] Samuel J. Gershman. “Origin of perseveration in the trade-off between reward and complexity”. Cognition. doi: 10.1016/j.cognition.2020.104394.

[40] Kentaro Katahira. “The statistical structures of reinforcement learning with asymmetric value updates”. Journal of Mathematical Psychology. doi: 10.1016/j.jmp.2018.09.002.

[41] Cooper D. Grossman, Bilal A. Bari, and Jeremiah Y. Cohen. “Serotonin neurons modulate learning rate through uncertainty”. Current Biology. doi: 10.1016/j.cub.2021.12.006.

[42] Kelly MJ Diederen and Wolfram Schultz. “Scaling prediction errors to reward variability benefits error-driven learning in humans”. Journal of neurophysiology.

[43] Nicola M. Grissom and Teresa M. Reyes. “Let’s call the whole thing off: evaluating gender and sex differences in executive function”. Neuropsychopharmacology. doi: 10.1038/s41386-018-0179-5.

[44] Cathy S. Chen et al. “Sex differences in learning from exploration”. eLife. doi: 10.7554/eLife.69748.

[45] Matthew R. Nassar et al. “Rational regulation of learning dynamics by pupil-linked arousal systems”. Nature Neuroscience. doi: 10.1038/nn.3130.

[46] Santiago Herce Castañón et al. “Human noise blindness drives suboptimal cognitive inference”. Nature communications.

[47] Junseok K Lee, Marion Rouault, and Valentin Wyart. “Adaptive tuning of human learning and choice variability to unexpected uncertainty”. Science Advances.

[48] Nicolas Schweighofer and Kenji Doya. “Meta-learning in reinforcement learning”. Neural Networks. doi: 10.1016/S0893-6080(02)00228-9.

[49] Shin Ishii, Wako Yoshida, and Junichiro Yoshimoto. “Control of exploitation-exploration meta-parameter in reinforcement learning”. Neural Networks. doi: 10.1016/S0893-6080(02)00056-4.

[50] Mehdi Khamassi et al. “Medial prefrontal cortex and the adaptive regulation of reinforcement learning parameters”. Progress in Brain Research. doi: 10.1016/B978-0-444-62604-2.00022-8.

[51] Massimo Silvetti et al. “Dorsal anterior cingulate-brainstem ensemble as a reinforcement meta-learner”. PLoS Computational Biology. doi: 10.1371/journal.pcbi.1006370.

[52] Dougal G. R. Tervo et al. “Behavioral variability through stochastic choice and its gating by anterior cingulate cortex”. Cell. doi: 10.1016/j.cell.2014.08.037.

[53] Matthew F. S. Rushworth and Timothy E. J. Behrens. “Choice, uncertainty and value in prefrontal and cingulate cortex”. Nature Neuroscience. doi: 10.1038/nn2066.

[54] James Webb et al. “Foraging animals use dynamic Bayesian updating to model meta-uncertainty in environment representations”. PLoS computational biology.

[55] Robert C. Wilson et al. “Humans use directed and random exploration to solve the explore-exploit dilemma”. Journal of Experimental Psychology: General. doi: 10.1037/a0038199.

[56] Vincent D. Costa, Andrew R. Mitz, and Bruno B. Averbeck. “Subcortical substrates of explore-exploit decisions in primates”. Neuron. doi: 10.1016/j.neuron.2019.05.017.

[57] Maria K. Eckstein et al. “Reinforcement learning and Bayesian inference provide complementary models for the unique advantage of adolescents in stochastic reversal”. Developmental Cognitive Neuroscience. doi: 10.1016/j.dcn.2022.101106.

[58] Alireza Soltani and Etienne Koechlin. “Computational models of adaptive behavior and prefrontal cortex”. Neuropsychopharmacology. doi: 10.1038/s41386-021-01123-1.

[59] Shiva Farashahi et al. “Metaplasticity as a Neural Substrate for Adaptive Learning and Choice under Uncertainty”. Neuron. doi: 10.1016/j.neuron.2017.03.044.

[60] Payam Piray and Nathaniel D Daw. “Computational processes of simultaneous learning of stochasticity and volatility in humans”. Nature communications.

[61] Joseph T. McGuire et al. “Functionally Dissociable Influences on Learning Rate in a Dynamic Environment”. Neuron. doi: 10.1016/j.neuron.2014.10.013.

[62] Marieke Jepma et al. “Catecholaminergic Regulation of Learning Rate in a Dynamic Environment”. PLoS Computational Biology. doi: 10.1371/journal.pcbi.1005171.

[63] Alessandro Piccin et al. “Orbitofrontal noradrenaline acts as an early gate for reversal learning”. Cell Reports.

[64] Stephanie M. Groman et al. “Orbitofrontal circuits control multiple reinforcement-learning processes”. Neuron. doi: 10.1016/j.neuron.2019.05.042.

[65] Juan Luis Romero-Sosa et al. “Neural coding of choice and outcome are modulated by uncertainty in orbitofrontal but not secondary motor cortex”. Nature Communications. doi: 10.1038/s41467-025-63866-5.

[66] Elise Payzan-LeNestour et al. “The Neural Representation of Unexpected Uncertainty during Value-Based Decision Making”. Neuron. doi: 10.1016/j.neuron.2013.04.037.

[67] Peter Dayan and Angela Jyu. “Uncertainty and Learning”. IETE Journal of Research. doi: 10.1080/03772063.2003.11416335.

[68] Peyman Khorsand and Alireza Soltani. “Optimal structure of metaplasticity for adaptive learning”. PLOS Computational Biology. doi: 10.1371/journal.pcbi.1005630.

[69] Charles Findling et al. “Computational noise in reward-guided learning drives behavioral variability in volatile environments”. Nature neuroscience.

[70] Maarten Speekenbrink and Emmanouil Konstantinidis. “Uncertainty and Exploration in a Restless Bandit Problem”. Topics in Cognitive Science. doi: 10.1111/tops.12145.

[71] Samuel J. Gershman. “Deconstructing the human algorithms for exploration”. Cognition. doi: 10.1016/j.cognition.2017.12.014.

[72] Michael Browning et al. “Anxious individuals have difficulty learning the causal statistics of aversive environments”. Nature Neuroscience. doi: 10.1038/nn.3961.

[73] Lucina Q. Uddin. “Cognitive and behavioural flexibility: neural mechanisms and clinical considerations”. Nature Reviews Neuroscience. doi: 10.1038/s41583-021-00428-w.

[74] Erdem Pulcu and Michael Browning. “The misestimation of uncertainty in affective disorders”. Trends in cognitive sciences.

[75] Rebecca P. Lawson, Christoph Mathys, and Geraint Rees. “Adults with autism overestimate the volatility of the sensory environment”. Nature Neuroscience. doi: 10.1038/nn.4615.

[76] Rebecca P. Lawson et al. “The computational, pharmacological, and physiological determinants of sensory learning under uncertainty”. Current Biology. doi: 10.1016/j.cub.2020.10.043.

